# Nucleolar-based *Dux* repression is essential for 2-cell stage exit

**DOI:** 10.1101/2021.11.11.468235

**Authors:** Sheila Q. Xie, Bryony J. Leeke, Chad Whidling, Ryan T. Wagner, Ferran Garcia-Llagostera, Paul Chammas, Nathan T-F. Cheung, Dirk Dormann, Michael T. McManus, Michelle Percharde

## Abstract

Upon fertilisation, the mammalian embryo must switch from dependence on maternal transcripts to transcribing its own genome, and in mice involves the transient upregulation of MERVL transposons and MERVL-driven genes at the 2-cell stage. The mechanisms and requirement for MERVL and 2-cell (2C) gene upregulation are poorly understood. Moreover, this MERVL-driven transcriptional program must be rapidly shut off to allow 2C exit and developmental progression. Here, we report that robust ribosomal RNA (rRNA) synthesis and nucleolar maturation are essential for exit from the 2C state. 2C-like cells and 2C embryos show similar immature nucleoli with altered structure and reduced rRNA output. We reveal that nucleolar disruption via blocking Pol I activity or preventing nucleolar phase separation enhances conversion to a 2C-like state in embryonic stem cells (ESCs) by detachment of the MERVL activator *Dux* from the nucleolar surface. In embryos, nucleolar disruption prevents proper *Dux* silencing and leads to 2-4 cell arrest. Our findings reveal an intriguing link between rRNA synthesis, nucleolar maturation and gene repression during early development.

## Introduction

Upon fertilisation, one of the earliest requirements for the development of a new organism is the formation of a totipotent zygote, which possesses the capacity to generate the entire embryo and all extra-embryonic structures. In mice, only the zygote and 2-cell stage embryo possess totipotency(Tarkowski 1959; Casser et al. 2017) with subsequent cleavages entailing a decrease in cellular plasticity as cells become specialised. Cells of the E4.5 epiblast, for example, are pluripotent, possessing the ability to generate all three germ layers of the embryo yet typically not extra-embryonic cell types(Rossant et al. 2003; Martinez Arias et al. 2013). Concurrent with the establishment of totipotency is the essential switch from reliance on maternal transcripts to activation of the embryo’s own genome, termed zygotic or embryonic genome activation (ZGA/EGA). Interestingly, ZGA and totipotency at the 2-cell stage have been linked to the rapid and transient activation of several families of transposable elements (TEs), most notably MERVL(Peaston et al. 2004; Svoboda et al. 2004; Macfarlan et al. 2012).

TEs have contributed a widespread and significant source of cis-regulatory information to mammalian genomes, providing transcription factor binding sites, enhancers, and promoter sequences(Kunarso et al. 2010; Chuong et al. 2013; Sundaram et al. 2014). Many 2-cell-specific and ZGA transcripts use MERVL LTR sequences as promoters, making the MERVL-dependent transcriptome an important component of ZGA(Macfarlan et al. 2011; Macfarlan et al. 2012). In humans, specific TEs from the HERVL family are also expressed upon EGA at the 4-8 cell stage(De Iaco et al. 2017; Hendrickson et al. 2017). Several studies suggest that correct MERVL regulation is functionally important during embryogenesis. MERVL depletion impairs developmental progression(Huang et al. 2017), while overexpression in embryonic stem cells (ESCs) confers expanded fate potential: the ability in chimeras to generate both embryonic and extra-embryonic lineages, similar to 2-cell blastomeres(Yang et al. 2020). However, the functional relevance of these TEs at ZGA, as well how and why they are swiftly repressed, is still poorly understood.

Understanding of the 2-cell stage and ZGA has been enhanced by the identification of a rare, transient population of cells within ESC cultures that share several epigenetic, metabolic, and transcriptomic features with 2-cell embryos, termed 2-cell (2C)-like cells(Macfarlan et al. 2012; Boskovic et al. 2014), marked by expression of a MERVL-GFP (2C-GFP) reporter. This tool recently led to the discovery of a Dux (DUX4 in human) as a potent MERVL/HERVL and 2C activator. Dux binding directly to 2C/MERVL promoters is sufficient to convert ESCs to a 2C-like fate, and in zygotes and early 2-cell embryos drives the expression of many early ZGA and 2C-specific genes(De Iaco et al. 2017; Hendrickson et al. 2017; Whiddon et al. 2017). Since then, several 2C-activators both upstream and downstream of *Dux* have been uncovered, including both transcriptional and post-transcriptional regulators(Choi et al. 2017; Guallar et al. 2018; Eckersley-Maslin et al. 2019; Hu et al. 2020).

Surprisingly, *Dux* knockout in embryos has overall mild effects, implying the existence of parallel and redundant mechanisms to activate MERVL and ZGA *in vivo*, which remain to be discovered(Chen and Zhang 2019; Guo et al. 2019; De Iaco et al. 2020; Bosnakovski et al. 2021). In contrast, the swift attenuation of *Dux* and MERVL expression for 2-cell stage exit is likely essential both *in vitro* and *in vivo*. Dux overexpression arrests embryos at the 2-4 cell stage(Guo et al. 2019), while prolonged *Dux* overexpression in ESCs causes DNA-damage and apoptosis(Olbrich et al. 2021). Similarly, DUX4 derepression in muscle cells causes the human disease, Facioscapulohumeral Muscular Dystrophy (FSHD), characterised by upregulation of DUX4 target genes, dsRNAs, TEs and apoptosis(Dixit et al. 2007; Geng et al. 2012; Shadle et al. 2017). Despite its importance, the mechanism for such rapid shutdown of *Dux* and MERVL gene expression at the late 2-cell stage is unclear.

Towards this, we recently reported a novel complex that is essential for *Dux* and MERVL/2C repression during early development, comprising the TE, LINE1 in association with Nucleolin (Ncl) and Kap1/Trim28 proteins(Percharde et al. 2018). LINE1 RNA in this complex is essential for proper *Dux* repression and its depletion induces the conversion of ESCs to the 2C-like state and causes 2-cell arrest in embryos(Percharde et al. 2018). At the same time, the discovery of Ncl as a *Dux* repressor implied an intriguing potential role for the nucleolus in 2-cell exit, which has not yet been explored.

Here, we investigated the impact of nucleolar dynamics and its link to *Dux* repression and 2-cell exit, using a new 2C-GFP reporter cell system and early mouse embryos. We find that 2C-like cells possess immature nucleoli with morphology akin to nucleolar precursor bodies (NPBs) that show reduced output and abrogated *Dux* repression compared to ESCs. Direct disruption of nucleolar structure and function by RNA Polymerase I inhibition (iPol I) or by perturbation of nucleolar liquid-liquid phase separation is sufficient to rapidly release *Dux* from perinucleolar regions, activate its expression, and convert ESCs into a 2C-like state. *In vivo*, short-term iPol I activates *Dux* and impairs developmental progression past the 2-4 cell stage. Our study reveals a direct link between rRNA transcription, nucleolar-based repression and cell fate during early mammalian development.

## Results

### The 2C-GFP/CD4 reporter enables rapid isolation of endogenous 2C-like cells

2-cell(2C)-like cells can be identified from within ESC cultures by expression of a stably-integrated fluorescent reporter (eg MERVL-GFP, 2C-GFP(Macfarlan et al. 2012; Ishiuchi et al. 2015)). These cells arise infrequently and transiently at a typical rate of less than 1-2%, making it challenging to perform large-scale or unbiased analyses in spontaneously-arising cells. Purification by flow cytometry assisted cell sorting (FACS) is laborious and slow, thus potentially perturbing biological processes(Binek et al. 2019). To perform 2C-like cell characterisation without flow sorting, we devised an improved strategy to allow FACS-free and rapid isolation of 2C-like cells. We generated ESCs stably harbouring a modified MERVL-GFP reporter, which induces expression of the extracellular portion of CD4 protein as well as GFP in the 2C state (2C-GFP/CD4+, Figure 1A). With this technique, naturally-arising 2C-like cells can be rapidly purified from ESC cultures by magnetic bead-based isolation with a typical purity of 55-85% after only 15 minutes, more than a 100-fold increase over the starting population (Figure 1B-C, Figure S1A-B). We confirmed that 2C-GFP/CD4+ (“2C-pos”) cells express markers of bona fide 2C-like cells, including high levels of MERVL and 2C-specific transcripts (Figure 1D, S1C). 2C-GFP/CD4+ cells display induction of MERVL Gag protein, together with loss of Oct4 protein and DAPI-dense chromocenters (Figure 1E-F), which are all previously-described features of 2C-like cells, and similar to 2-cell embryos (Macfarlan et al. 2012; Ishiuchi et al. 2015; Percharde et al. 2018). Thus, 2C-GFP/CD4+ cells faithfully capitulate the 2C-like state.

**Figure 1.**
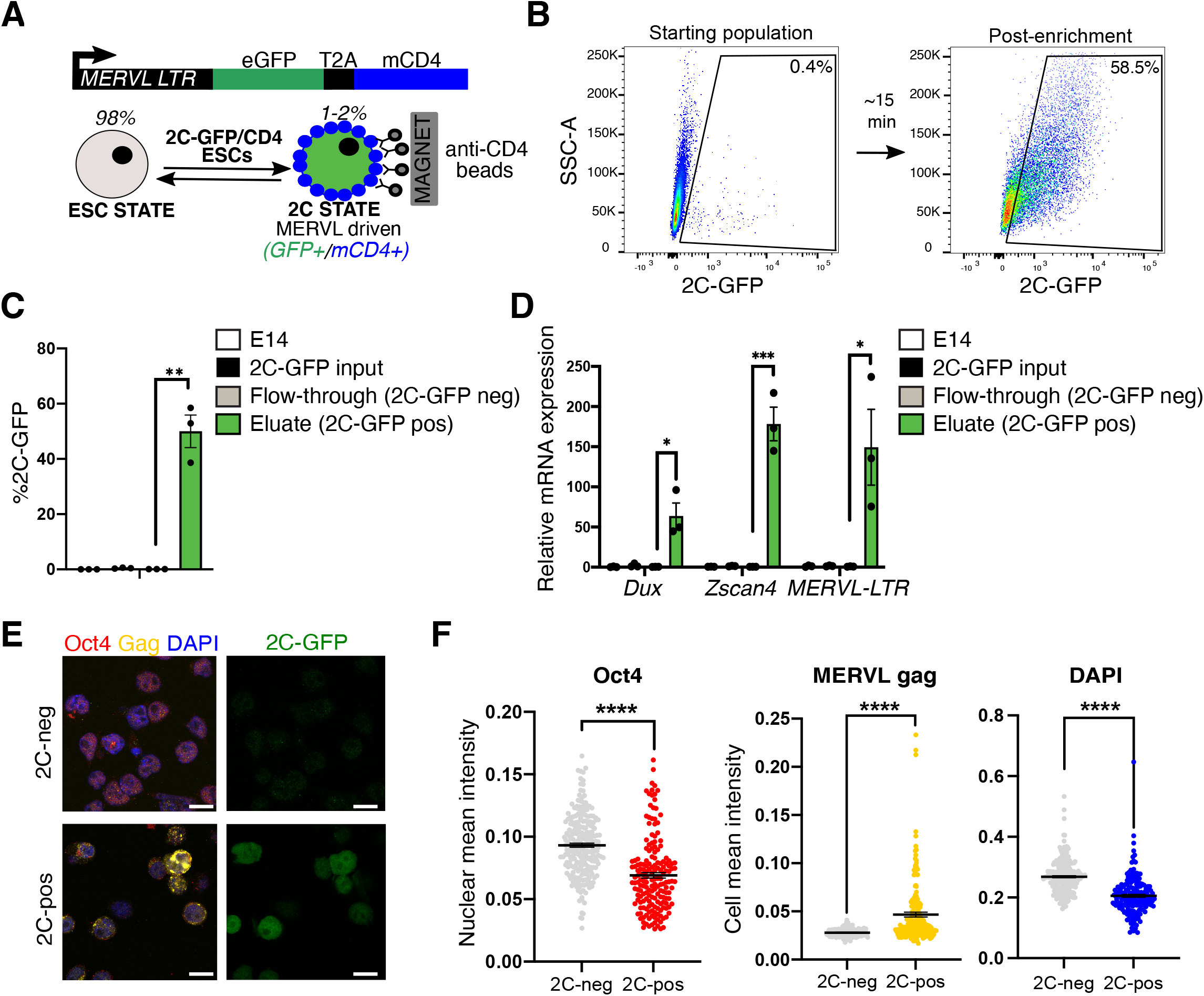
A new reporter cell line for purification of 2C-like cells. (A) Reporter design: a previous MERVL-GFP reporter(Ishiuchi et al. 2015) is modified to contain the extracellular portion of the CD4 antigen downstream of GFP and a T2A cleavage element, allowing rapid 2C-like cell purification by anti-CD4 beads. (B) Representative flow cytometry plot depicting proportion of typical 2C-GFP+ (2C-pos) cells before (left) and after (right) CD4-based 2C enrichment. (C) Percent recovery of 2C-GFP positive (pos) cells after CD4-based purification, comparing CD4+ cells (eluate) and CD4-cells (flow-through). Data are mean +/− s.e.m of 3 experiments. (D) qRT-PCR validation of high levels of 2C-specific genes and TEs in the CD4+ eluate compared to CD4-fraction and the starting population. Data are mean +/− s.e.m of 3 experiments. (E) Representative confocal images and (F) quantification of levels of Oct4 and MERVL gag proteins and DAPI in 2C-pos versus 2C-neg cells following CD4-based purification. Scale bar, 20 μm All P values represent two-tailed, unpaired Student’s t-test, with Welch correction for uneven variance where relevant.

### 2C-like cell nucleoli resemble NPBs and exhibit reduced nucleolar function

We previously discovered that a ribonucleoprotein complex comprising LINE1 RNA, together with Nucleolin (Ncl) and Kap1, is essential for both ribosomal RNA (rRNA) expression as well as 2-cell exit(Percharde et al. 2018). Since Ncl and rRNA are both well-known nucleolar components, we investigated whether the 2C-like state is associated with changes to nucleoli. 2C-positive and negative cells were isolated following CD4 enrichment and examined by confocal microscopy (Figure 2A). Interestingly, we found that 2C-like cells possess a distinct nucleolar morphology, with a rounded, ring-like structure (Figure 2B-C). We next tested whether nucleolar morphological changes in 2C-like cells might also be accompanied by changes to RNA Polymerase I activity and nucleolar function. Nucleoli are the site of RNA Polymerase I-driven ribosomal RNA (rRNA) synthesis, processing, and ribosomal assembly. rRNA makes up over 70% of cellular RNA, which is tightly co-ordinated with *Rpl/Rps* RNA expression and protein synthesis(Laferte et al. 2006; Percharde et al. 2017). We measured production of nascent RNA and protein in the 2C-like versus ESC state using Click-iT assays, where a pulse of nucleotides or amino acid analogues is given to cells that are then fluorescently labelled post fixation for quantification (Figure 2A). We discovered a significant reduction in translation in 2C-like cells (Figure 2D) as well as reduced nascent RNA synthesis – the majority of which comprises nucleolar rRNA (Figure 2E, S2A, and inset). To confirm that these changes are not an artefact of CD4-based enrichment, nascent transcription and translation rates were profiled in unsorted, bulk 2C-GFP reporter ESCs(Percharde et al. 2018). In agreement, spontaneously-arising 2C-like cells exhibit reductions in nucleolar function in contrast to neighbouring ESCs (Figure 2F).

**Figure 2.**
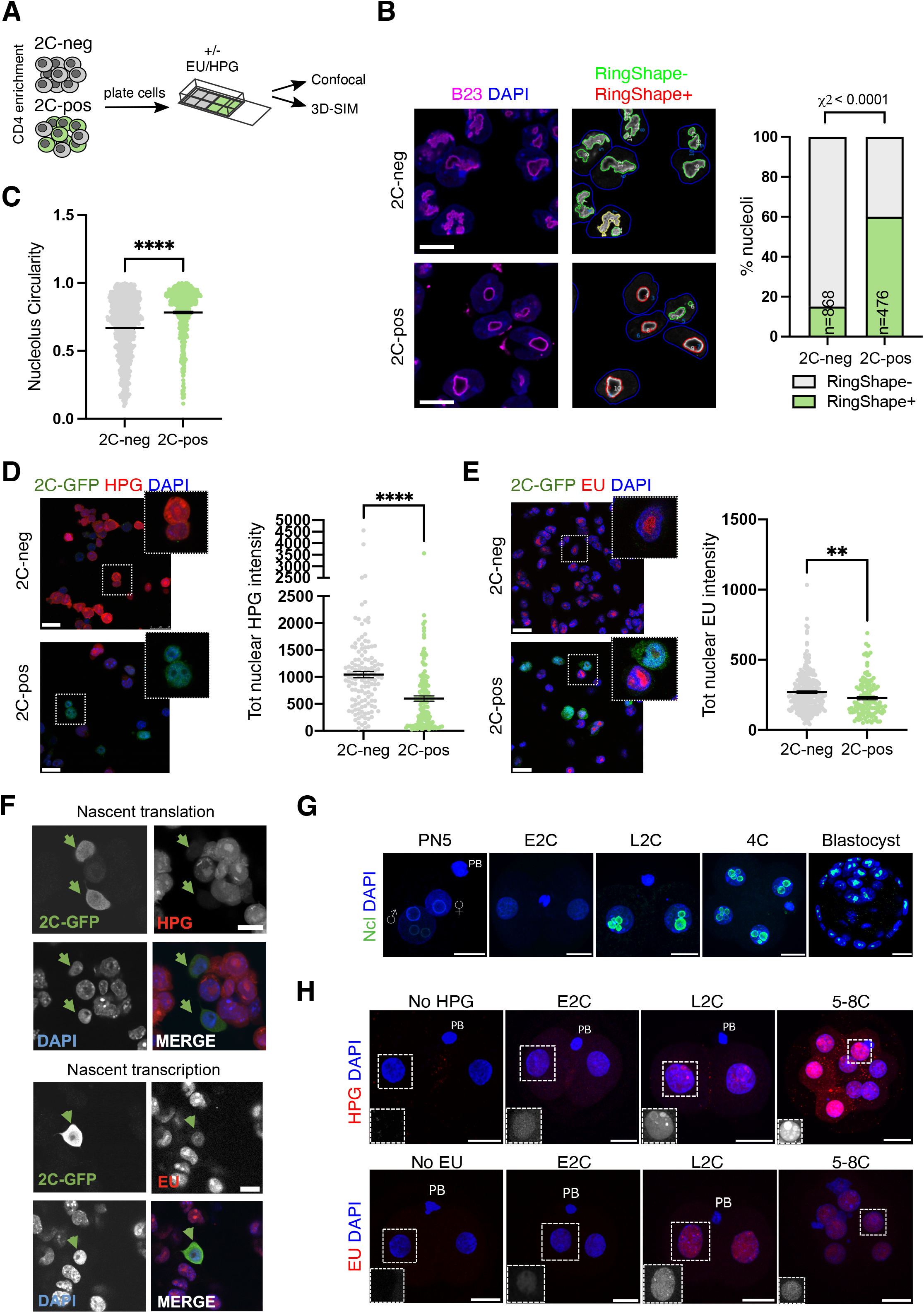
2C-like cells and embryos have altered nucleolar morphology and function. (A) Experimental set-up for 2C-like cell profiling: following CD4-based enrichment, 2C-neg/pos populations are plated into Matrigel-coated chambers for a minimum of 1h before the indicated downstream applications. (B) Immunofluorescence images and quantification (RingShape+, CellProfiler) appearance in 2C-pos/neg cells, revealing that 2C-like cell nucleoli, stained by the nucleolar marker B23 (Npm1), have rounded, ring-like morphology, n, number of nucleoli scored. Scale bar, 20 μm. (C) Nucleolar circularity is significantly increased in 2C-like cells. Very small nucleoli (area <100 pixels) were filtered out as can typically generate unreliable measurements (see methods). (D-E) Immunofluorescence images and quantification of (D) nascent translation and (E) nascent transcription rates in 2C-pos versus 2C-neg cells via HPG or EU Click-iT incorporation experiments, respectively. Scale bar, 25 μm. (F) Confirmation of reduced transcription and translation in 2C-like (2C-GFP+) cells within unsorted populations, using an independent 2C-GFP cell line(Percharde et al. 2018). Scale bar, 10 μm. (G) Nucleolar (Nucleolin, Ncl) staining in in vitro cultured embryos, showing the emergence of Ncl+ nucleoli at the late 2-cell stage (L2C). PN5, zygote PN5 stage, E2C, early 2-cell stage; PB, polar body. Scale bar, 20 μm (H) Analysis of nascent transcription/translation in embryos by EU/HPG assays. Scale bar, 20 μm. Insets show EU/HPG staining alone (grayscale) in a representative blastomere from each image. P values represent (B) Chi-squared test and (C, D-E) two-tailed Student’s t-test, with Welch’s correction for uneven variance where relevant.

Subsequently, we investigated whether these changes are reflected at the 2-cell stage *in vivo*. Following fertilisation, 1-2 cell embryos possess immature nucleolar precursor bodies (NPBs) - largely uncharacterised structures that are initially transcriptionally silent and lacking distinct compartments(Flechon and Kopecny 1998). In contrast, mature nucleoli contain 3 sub-compartments: a fibrillar centre surrounded by a dense fibrillar component, which itself is surrounded by a granular component. In contrast to mature nucleoli, embryo NPBs are morphologically similar to 2C-like nucleoli (Figure 2B, 2G). Coincident with the increasing initiation of rRNA transcription, mature nucleoli only gradually form from NPBs at the late 2-cell stage onwards(Kyogoku et al. 2014; Borsos and Torres-Padilla 2016). We analysed nucleolar function in embryos with nascent transcription/translation assays, which demonstrated dynamic rates of biosynthesis during the 2-cell stage. Early 2-cell (E2C) embryos exhibit low levels of nascent RNA synthesis but also nucleolar translation, which rapidly increases by the late 2C (L2C) stage and upon 2-cell exit (Figure 2H). At the same time, Ncl protein only becomes readily detectable surrounding nucleoli in L2C embryos onwards (Figure 2G), at the time when MERVL and the 2-cell program is being shut down. We conclude that the 2-cell stage *in vitro* and *in vivo* is characterised by significantly reduced nucleolar function and morphologically distinct nucleolar structure.

### Nucleolar disruption induces conversion to the 2C-like state

The observed nucleolar remodelling upon 2-cell stage exit lead us to ask whether alterations to nucleolar structure and function might be drivers of the 2C-like state. We took advantage of two different small molecules to inhibit Pol I and rRNA synthesis, CX-5461 - which blocks recruitment of the Pol I initiation factor SL1 to rDNA(Bywater et al. 2012; Haddach et al. 2012), and BMH-21, which triggers rapid Pol I degradation(Peltonen et al. 2014). We found that nucleolar disruption by mild or partial inhibition of rRNA synthesis (Pol I inhibition, iPol I) is detectable by 2h (Figure S3A) and by 4h induces morphological nucleolar remodelling, generating singular ring-like structures in ESCs resembling 2C-like nucleoli and embryo NPBs (Figure 3B, Figure 2G). Importantly, the structures observed following this milder inhibition are distinct from the nucleolar cap-like structures seen upon more extreme nucleolar stress, where fibrillar proteins such as Fibrillarin (Fbl) or UBF aggregate at the nucleolar periphery, or from complete dissolution of nucleolar proteins into the nucleoplasm (Figure S3B)(Shav-Tal et al. 2005; Ide et al. 2020). Moreover, we did not detect gross changes to Ncl or Fbl protein abundance upon iPol I (Figure S3C). Strikingly, we found that following nucleolar reprogramming, iPol I causes a significant increase to 20% of 2C-like cells within ESC cultures (Figure 3C). In agreement, iPol I induces high expression of 2C-specific genes and MERVL transposons (Figure 3D) along with 2C proteins, Zscan4 and MERVL gag in ESCs (Figure 3E). Thus, nucleolar disruption produces 2C-like nucleoli and moreover is sufficient to reprogram ESCs into the 2C-like state.

**Figure 3.**
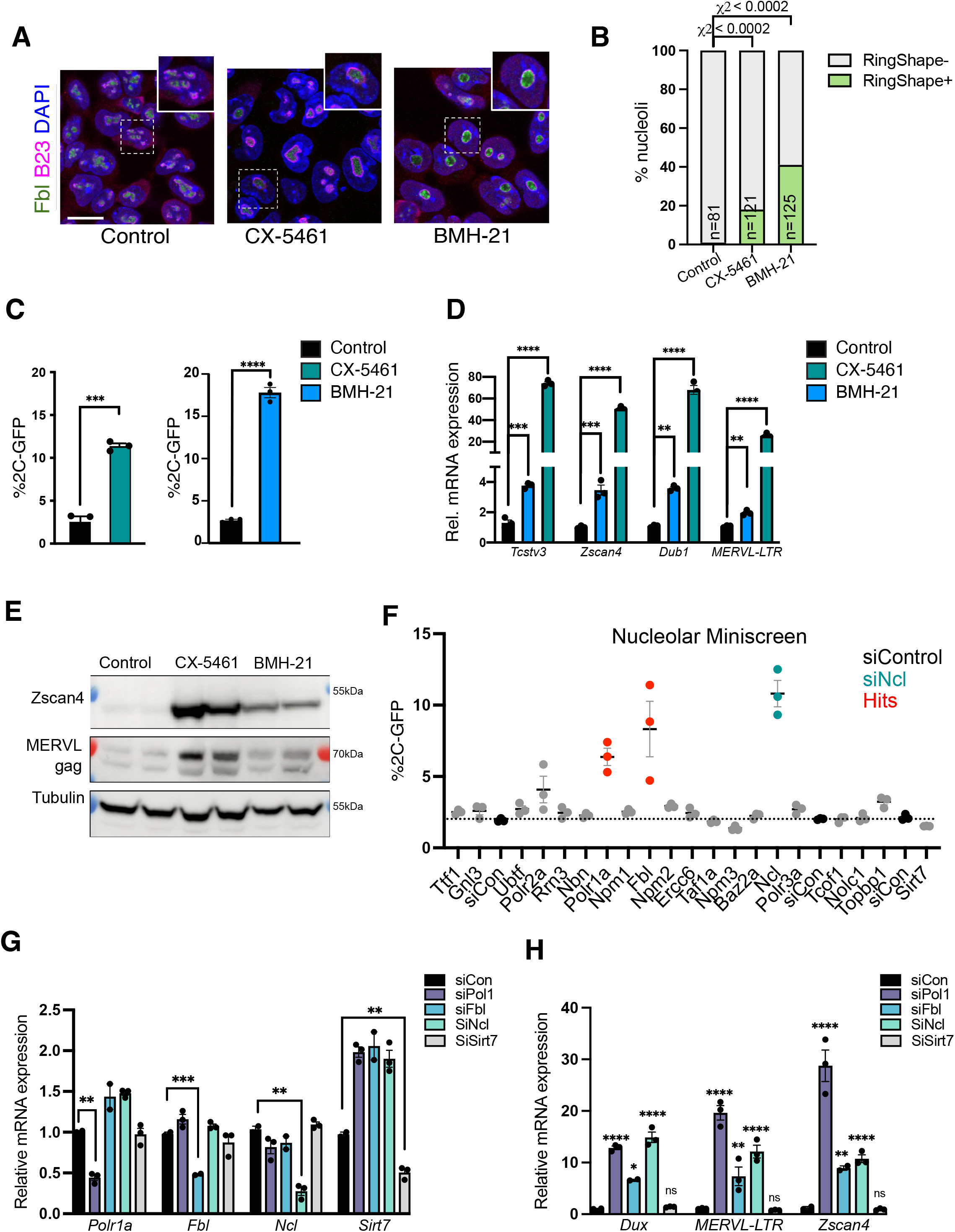
Nucleolar disruption induces the 2C-like state. (A) Representative immunofluorescence images following staining for nucleolar markers (Fibrillarin, Fbl) and B23 4h after RNA Pol I inhibition (iPol I), with either CX-5461 or BMH-21. Scale bar, 20 μm. (B) Quantification of the percentage of ring-like (RingShape+) nucleoli 4h after iPol I. P values, Chi-squared test; n, number of nucleoli. (C) Percentage of 2C-GFP+ cells following overnight (16-24h) treatment with 0.25 μM iPol I, data are mean +/− s.e.m, n=3 biological replicates representative of 3+ experiments. (D) qRT-PCR analysis of 2C-specific genes and TEs following iPol I as in (C), with P values in (C-D) representing 2-tailed t-test with two-stage multiple comparisons correction. (E) Western blots showing upregulation of 2C-specific proteins, Zscan4 and MERVL gag, after 16-24h iPol I in ESCs. Replicates from two experiments are shown. (F) Flow cytometry analysis of % 2C-GFP+ cells following siRNA knockdown of the indicated factors. Red samples indicate a Z-score of >1, with KD of Ncl shown as a positive control (teal). Data are mean +/− s.e.m of 3 biological replicates, representative of two repeats of the screen. (G) Validation by qRT-PCR of siRNA-mediated knockdown of the indicated factors and (H) upregulation of 2C-specific genes, showing mean +/− s.e.m of n=2-3 biological replicates, representative of 2 experiments. P values, 2-way ANOVA followed by Dunnett’s multiple comparisons test.

### Nucleolar proteins driving rRNA synthesis and processing are essential for *Dux* and 2C-repression

These results support the hypothesis that the development of functionally mature nucleoli may play a role in the repression of the 2-cell transcriptional program and for exit from the 2-cell stage. To investigate this, we asked which nucleolar proteins are most important for repression of the 2C-like state in ESCs and performed an siRNA miniscreen for nucleolar components in 2C-GFP ESCs. Similar to the effects of Ncl loss, depletion of Pol I and Fibrillarin (Fbl) also cause a notable increase in 2C-like cells (Figure 3F, S3D), while knockdown (KD) of other nucleolar proteins has limited effect. Conversely, KD of Npm3, a negative regulator of ribosome biogenesis(Huang et al. 2005), leads to a small but consistent reduction in 2C-GFP+ cells (Figure 3F). Confirming these results, siRNAs against Pol I, Fbl and Ncl all induce high levels of 2C-specific genes and transposons (Figure 3G-H), indicating that these factors are necessary for repression of the 2C-like state. Interestingly, RNA Pol I, Ncl and Fbl are all known to be critical for rRNA synthesis and/or processing(Ginisty et al. 1998; Yao et al. 2019; Ide et al. 2020). Collectively, these data reveal an intriguing link between rRNA synthesis and 2C-repression to maintain ESC identity.

Next, we asked how rRNA synthesis and nucleolar function is mechanistically linked to repression of the 2C-like state. We focused on *Dux*, which is a potent 2C activator and is upregulated upon nucleolar protein knockdown (Figure 3H). We performed time-course experiments of acute iPol I treatment followed by qRT-PCR and RNA-seq (Figure 4A, S4A), and found that *Dux* is significantly induced as early as 4h following nucleolar disruption (Figure 4A, S4B). By 8h, *Dux* targets are highly upregulated amongst all significantly altered genes following iPol I (Figure 4B). Moreover, transcriptomic profiling of 2C-specific genes(Macfarlan et al. 2012; Percharde et al. 2018) demonstrated that MERVL and the 2C program is widely upregulated following nucleolar disruption (Figure 4C, S4B-C). Next, to test whether 2C gene induction is dependent on Dux, we performed iPol I experiments with *Dux* siRNAs. We confirmed that 2C genes are specifically upregulated by 8h iPol I, which is prevented upon *Dux* depletion (Figure 4D-E). We subsequently investigated whether nucleolar disruption can also prevent *Dux* silencing in embryos (Figure 4F), which normally occurs rapidly as embryos transit to the late 2-cell stage (Figure 4G and(De Iaco et al. 2017; Hendrickson et al. 2017)). We found that iPol I treatment in mid-2C embryos leads to significant rRNA reduction and concomitant *Dux* activation in L2C-4C embryos (Figure 4G-H). We observed slightly different kinetics with the two inhibitors, with BMH-21 causing more rapid activation of *Dux* in embryos than CX-5461 (Figure 4G-H), similar to in ESCs (Figure 4A). Finally, nucleolar disruption leads to an inability to progress beyond 2-4 cell stage, in contrast to control embryos (Figure 4I, S4D). Together, these results indicate that nucleolar disruption rapidly leads to *Dux* de-repression and promotion of the 2-cell state both in ESCs and in embryos.

**Figure 4.**
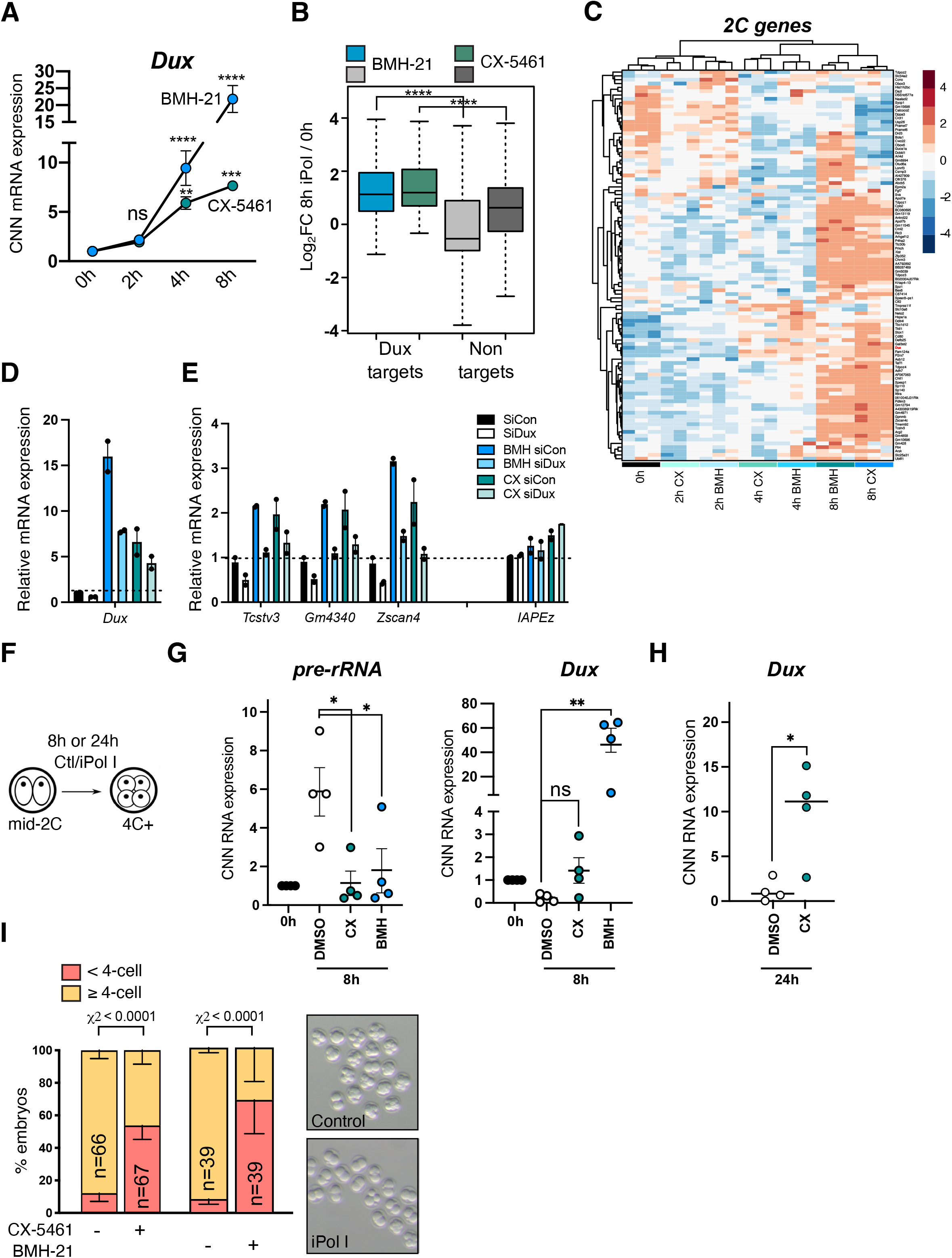
Nucleolar disruption causes *Dux* reactivation in ESCs and embryos. (A) Cell-number normalised (CNN) qRT-PCR time course analysis of *Dux* upregulation following iPol I. Data are analysed as CNN to exclude potential global effects of iPol I on transcription; however the same results are seen with *Rpl7/H2A* normalisation. Data are mean +/− s.e.m, n=3 biological replicates. P values, two-way ANOVA and Šídák multiple comparisons test. (B) Boxplot of log2-fold change values for n=99 *Dux* target genes(Percharde et al. 2018) versus significantly altered non-targets (FDR <0.05, n=8057: CX-5461, n=13830: BMH-21) following 8h iPol I. P values, two-sided Wilcoxon rank-sum test. (C) Heatmap of 2C-specific genes(Macfarlan et al. 2012) showing gradual upregulation following iPol I. Samples are grouped by unsupervised hierarchical clustering. (D-E) Expression of (D) *Dux* and (E) 2C-specific genes or a negative control (IAPEz) in iPol I rescue experiments with control or *Dux* siRNAs. Data are mean +/− s.e.m of two independent cell lines, representative of 2 experiments. (F) Schematic for embryo iPol I inhibitor experiments with 1 μM BMH-21 or CX-5461. (G) CNN qRT-PCR expression data following 8h iPol I in mid 2-cell embryos showing inhibited *Dux* repression. Data are mean +/− s.e.m, n=4 experiments with equal numbers of embryos, with levels at 0 h set to 1 in each experiment. P values, 1-way ANOVA with Dunnett multiple comparisons correction. (H) CNN qRT-PCR expression data showing *Dux* upregulation after 24h 1 μM CX-5461. Data are mean +/− s.e.m n=4 experiments. P values, Welch’s two-tailed t-test. (I) Embryo progression rates following 24h iPol I treatment in n=4 experiments (CX-5461) and n=2 experiments (BMH-21). P values, Chi-squared test, n=number of embryos.

### *Dux* is repressed in perinucleolar chromatin

Prolonged treatment or high doses of drugs that perturb rRNA synthesis or cause rDNA damage is known to activate nucleolar stress. In this process, disruption of nucleolar integrity releases ribosomal proteins into the nucleoplasm to bind MDM2, leading to p53 stabilisation and activation followed by downstream effects such as cell cycle arrest(Rubbi and Milner 2003; James et al. 2014). To understand how nucleolar disruption is linked to *Dux* activation, we first tested whether this is dependent on nucleolar stress. Although iPol I does not induce typical markers of nucleolar stress (Figure S3B), levels of total and activated, phospho-p53 are increased upon iPol I, similar to the effect of the Topoisomerase II inhibitor, etoposide, used as a positive control (Figure S5A). Interestingly, etoposide treatment also increases the proportion of 2C-GFP+ cells in culture (Figure S5B), suggesting that p53 activation can activate the 2C-like state. Indeed, a recent study reported DNA damage-dependent activation of Dux/DUX4 and the 2C-like state via p53(Grow et al. 2021). However, we did not detect any increase in phospho-p53 in endogenously arising 2C-like cells (Figure S5C). Furthermore, iPol I treatment is still able to cause *Dux* activation in the absence of p53 (Figure S5D). Thus, although p53 activation is sufficient to induce the 2C-like state upon DNA damage(Grow et al. 2021), it is not necessary for *Dux* de-repression upon nucleolar disruption.

In ESCs but not 2C-like cells, we reported that *Dux* genes localise to perinucleolar regions with unknown functional relevance(Percharde et al. 2018). Mature nucleoli are surrounded by a shell of chromatin that is enriched for repressive histone marks(Nemeth et al. 2010; Lu et al. 2020) and is lowly transcribed(Quinodoz et al. 2018). We therefore reasoned that disruption of nucleolar function and morphology might lead to *Dux* upregulation by preventing its repression at the nucleolar periphery. To observe perinucleolar chromatin in more detail, we performed 3D super-resolution Structured Illumination Microscopy (3D-SIM) of DAPI staining in ESCs versus 2C-like cells. These experiments revealed a reduction in perinucleolar chromatin fibres in the 2C state (Figure 5A, orange arrows), alongside a previously-reported loss of chromocenters(Ishiuchi et al. 2015). Reduced nucleolar DNA association is moreover replicated upon BMH-21 and CX-5461 treatment (Figure 5B), suggesting that iPol I perturbs nucleolar-associated chromatin. We next examined whether nucleolar disruption alters the localisation of the *Dux* gene locus, focusing on acute inhibition to determine direct effects of iPol I. DNA FISH confirmed that *Dux* is frequently associated with perinucleolar regions in ESCs, and moreover revealed significant movement away from nucleoli to the nucleoplasm by 4h of either CX-5461 or BMH-21 (Figure 5C-D). Using 3D nuclear segmentation and analysis of *Dux* distance to nuclear compartments, we confirmed these findings and found that movement away from the nucleolus starts from 2h CX-5461 and robustly at 4h. In contrast, there is no change in *Dux* distance from the lamina (Figure S6A-C). Thus, it is only nucleolar-localised *Dux* alleles that are affected by iPol I. *Dux* movement is closely linked to reactivation of *Dux* (Figure 4A), with single-cell analysis of *Dux* transcription demonstrating that maximal nascent expression is reached by 4h iPol I (Figure S6D). Importantly, nascent *Dux* expression occurs only in the nucleoplasmic or lamina compartments, in strong agreement with the repressive nature of the nucleolus (Figure S6E).

**Figure 5.**
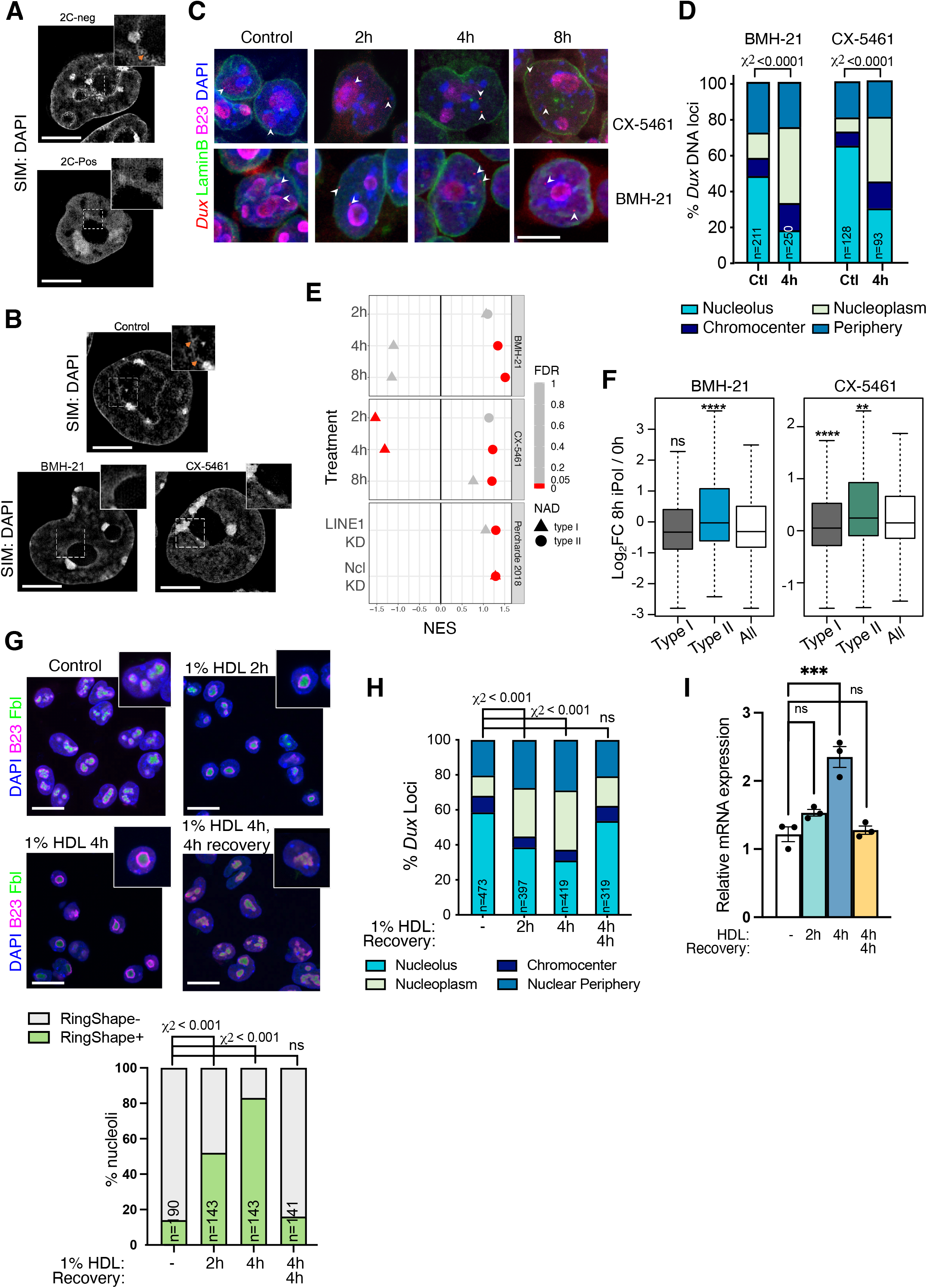
Nucleolar disruption induces *Dux* relocalisation and activation. (A-B) Example images of chromatin distribution as marked by DAPI staining in 3D-SIM imaging experiments in (A) 2C-pos versus 2C-neg cells and (B) in ESCs upon 8h iPol I. 2C-neg cells and (B) control but not iPol I ESCs have nucleolar chromatin fibres, visible as a roughened nucleolar border (orange arrows, inset). Scale bar, 5 μm. (C) Representative immuno-DNA FISH images at the indicated timepoints of iPol I for *Dux* alleles (red) compared to nucleolar (B23, magenta) or nuclear lamina (LaminB, green) compartments. Scale bar, 10 μm. (D) Quantification of *Dux* localisation at 4h iPol I showing movement away from the nucleolus, P values, Chi-squared test. N, number of loci (E) Dotplot of GSEA enrichment scores (NES) and significance (FDR) for Type I or Type II NADs using expression data following iPol I, or following LINE1/Ncl KD (Percharde et al. 2018). (F) Boxplot of log2-fold change values for Type I NADs (n=1565) or Type II NADs (n=371) versus all genes at 8h iPol I. P values, two-sided Wilcoxon rank-sum test, comparing Type I/II NADs to all genes. (G) Immunofluorescence for nucleolar markers B23 and Fbl after the indicated times of incubation with 1% 1,6-hexanediol (HDL), with or without washout and recovery in normal media, and (below) quantification of the percentage of RingShape+ nucleoli (n), scale bar, 20 μm. P values, Chi-squared test with Bonferroni adjustment for multiple comparisons. (H) Scoring of *Dux* loci nuclear positioning following HDL treatments from *Dux* immuno-FISH experiments; n, number of loci from two FISH experiments. P values, Chi-squared test, with Bonferroni adjustment for multiple comparisons. (I) Expression of *Dux* by qRT-PCR following HDL treatment, data are mean +/− s.e.m for n=3 biological replicates, representative of two independent experiments. P values, one-way ANOVA with Dunnett correction for multiple comparisons.

Nucleolar-associated DNA regions (NADs) have been previously identified by isolation and sequencing of DNA associated with purified nucleoli (NAD-seq(Nemeth et al. 2010; Vertii et al. 2019)), which has confirmed the generally repressive nature of nucleolar chromatin(Lu et al. 2020). Using NAD annotations generated from ESCs(Bizhanova et al. 2020), we asked whether the expression of other nucleolar-associated genes is altered following nucleolar disruption. We looked at Type I NADs, which overlap constitutively lamina associated domains (cLADs) and are considered to comprise constitutive heterochromatin, and Type II NADs, which do not overlap LADs in multiple cell types(Peric-Hupkes et al. 2010). GSEA analysis revealed that Type II NAD genes are particularly sensitive to nucleolar disruption, and are significantly upregulated from 4h iPol I compared to all genes, in contrast to Type I NADs (Figure 5E-F). This is not an isolated effect of inhibitor treatment, as knockdown of Ncl or LINE1, both important for nucleolar function(Percharde et al. 2018; Lu et al. 2020), also lead to NAD and NAD/LAD gene upregulation (Figure 5E). Together, these results reveal that nucleolar association of *Dux* is closely tied to its repression, and suggest that iPol I induces global disruption of nucleolar chromatin organisation and gene expression.

### Phase-separated nucleolar integrity is required for *Dux* repression at nucleoli

Lastly, we sought to determine the link between disrupted rRNA synthesis and *Dux* loci release and derepression. The membrane-less nucleolus is held together by liquid-liquid phase separation (LLPS), which is driven by the association of rDNA with nucleolar proteins and moreover dependent on continual rRNA synthesis(Feric et al. 2016; Yao et al. 2019; Ide et al. 2020). We hypothesized that the disruption of rRNA synthesis may inhibit nucleolar integrity and LLPS, thus allowing the release of associated DNA regions such as *Dux*. To test this, we used 1,6-hexanediol (HDL), an aliphatic alcohol used to disrupt liquid-like condensates(Ribbeck and Gorlich 2002). Short-term treatment with 1% HDL – a dose notably lower than typically used to disrupt non-nucleolar compartments(Vertii et al. 2019) – is sufficient to alter nucleolar morphology, resembling 2C-like cells or iPol I treatment (Figure 5G). Disruption of phase separation remarkably releases *Dux* loci after only 2h HDL (Figure 5H), and moreover leads to significant *Dux* upregulation by 4h (Figure 5I). Furthermore, we found this to be highly dynamic, with nucleolar morphology, *Dux* localisation and transcriptional repression all returning to normal after 4h washout (Figure 5I). These results suggest that *Dux* localisation and repression is maintained in the nucleolus through LLPS. Taken together, our data show that nucleolar disruption by several means causes *Dux* reactivation, initiation of 2C/MERVL gene transcription and conversion back to the 2C-like state. Overall these findings point to a novel requirement for rRNA biogenesis, nucleolar maturation and nucleolar-based repression for correct cell identity during the earliest stages of embryo development.

## Discussion

Major ZGA is an essential process occurring at the 2-cell stage of early mouse embryogenesis, which entails rapid activation of zygotic RNAs required for subsequent development. This includes a significant number of transcripts driven by the TE, MERVL, which unlike other ZGA transcripts are rapidly downregulated upon 2-cell exit. These dynamics swiftly follow the rapid repression of the MERVL activator, Dux. Sustained Dux expression in 2-cell embryos is poorly tolerated and moreover promotes persistence of the 2C program and impedes development(Percharde et al. 2018; Guo et al. 2019). Thus, timely Dux repression is essential, yet the mechanisms for this process are poorly understood.

Here, we reveal that high rRNA synthesis and nucleolar maturation from inactive NPBs to be an essential driver of *Dux* repression in embryos and 2C-like cells. The absence of pluripotency proteins such as Oct4 is a well-known feature of the 2C-like state(Macfarlan et al. 2012). Here we place this finding within the context of suppression of both rRNA transcription and global translation and reveal that the 2C-like state is characterised by significantly reduced nucleolar function, akin to NPBs in 2-cell embryos. NPBs are unique structures that in 1-2 cell embryos lack distinct compartments and exhibit low rRNA synthesis(Flechon and Kopecny 1998; Borsos and Torres-Padilla 2016). Nucleolar maturation occurs with the resumption of transcription, and is essential to generate high levels of rRNA and promote ribosomal assembly to fuel embryonic growth. However, it is becoming clearer that nucleoli also possess other roles in development. NPBs are essential for early centromeric chromatin organisation, which localises to the surface of NPBs(Zuccotti et al. 2005; Fulka and Langerova 2014) and nucleolus removal in oocytes causes 2-cell arrest(Ogushi et al. 2008). Intriguingly, reprogramming to totipotency by somatic cell nuclear transfer (SCNT) generates NPBs after only 3h(Martin et al. 2006), highlighting a link between totipotency and nucleolar biology. Here, we additionally show that 2C-like cells have NPB-like nucleoli with reduced chromatin association. We propose that nucleolar maturation and full nucleolar function is critical for *Dux* recruitment to the nucleolar periphery for its repression, which is in turn essential for 2-cell exit. Conversely, mild inhibition of Pol I is sufficient to rapidly release *Dux* from nucleolar chromatin and to activate its expression (Figure 6). In this way, we hypothesise that ZGA itself provides the mechanism to shut down the 2C program, in a feedback loop whereby high levels of rRNA synthesis promote nucleolar maturation that can then silence Dux.

**Figure 6.**
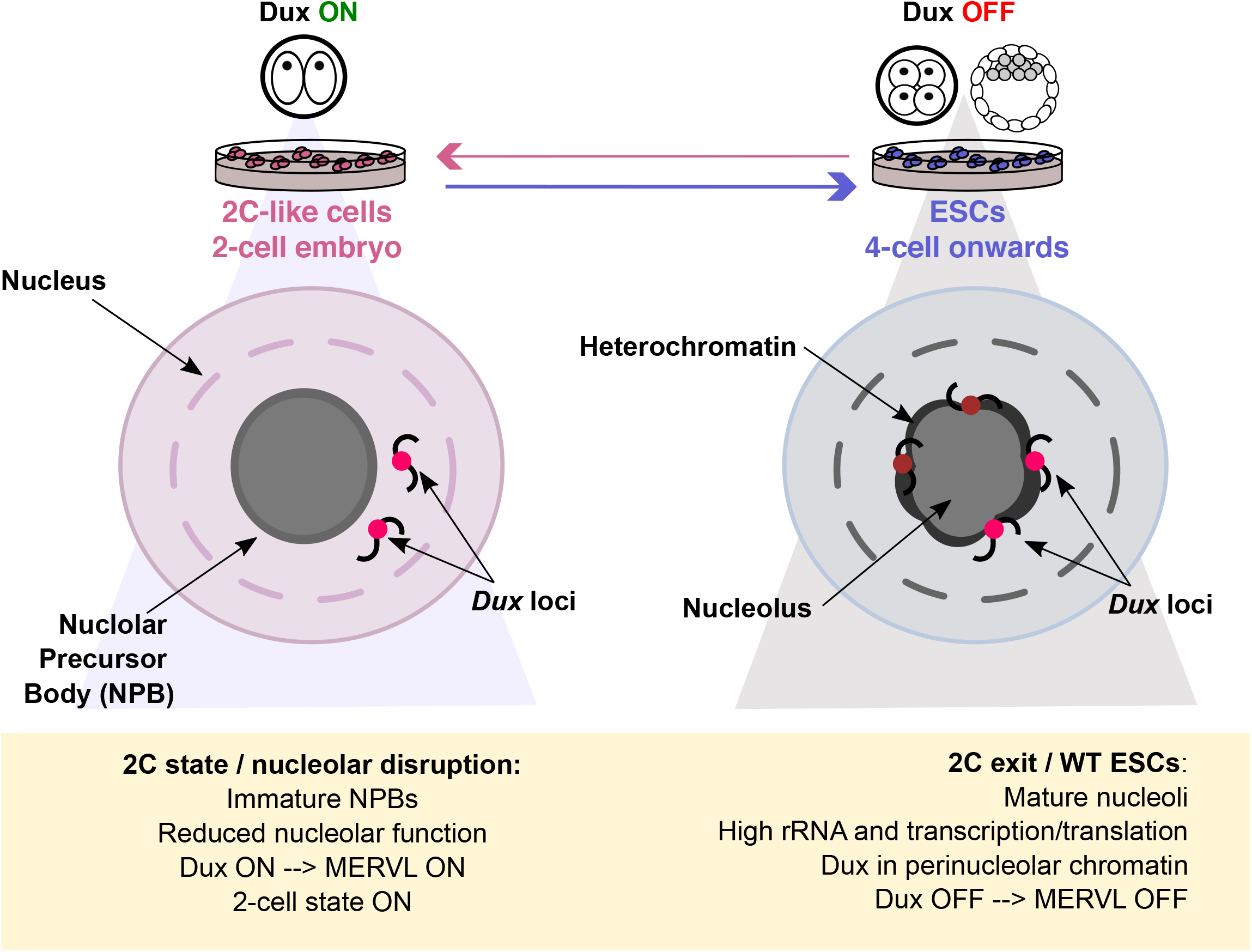
A model for nucleolar-based Dux and 2C-state repression. Nucleolar maturation allows for Dux repression and 2-cell exit. In early embryos and 2C-like cells, NPBs have altered morphology, reduced function and reduced chromatin association. We propose this provides a permissive environment for *Dux* and subsequent 2C/MERVL expression. In mature nucleoli with high rRNA output, *Dux* is recruited to perinucleolar chromatin and is repressed. Disruption of nucleolar integrity via iPol I or inhibition of nucleolar phase separation releases *Dux* and leads to its derepression.

Importantly, this mechanism of Dux regulation appears separate from nucleolar stress-mediated p53 activation, which is capable of directly inducing *Dux*(Grow et al. 2021). We find that iPol I ESCs or 2C-like cells do not display typical nucleolar stress markers and iPol I ESCs still activate *Dux* in the absence of p53. Instead, our data agree with previous nucleolar positioning studies, which demonstrated in yeast that an ectopic rDNA repeat can silence its chromosomal region(Zuccotti et al. 2005), and that 5S rDNA sequences are sufficient to induce nucleolar association and silencing of a reporter gene in ESCs(Fedoriw et al. 2012). Indeed, NAD-seq data(Bizhanova et al. 2020) indicate that *Dux* is located within a NAD (Figure S7). Interestingly, *D4Z4* repeats containing human *DUX4* have also been proposed to reside within a NAD(Nemeth et al. 2010) and bound by NCL(Gabellini et al. 2002). Studying nucleolar regulation of *DUX4* will allow us to understand if failure of a similar mechanism may contribute to DUX4 de-repression in the disease, FSHD.

Our data raise the question of how rRNA transcription is tied to *Dux* nucleolar association and repression. The nucleolus is self-organised into its three subdomains by phase separation, driven by the interaction between nucleolar proteins and rDNA(Feric et al. 2016; Yao et al. 2019), and nucleated by rRNA(Falahati et al. 2016). Indeed, purified Fbl and B23 (Npm1) can separate into distinct layers and recapitulate the dense fibrillar component and granular component in solution(Feric et al. 2016). Importantly, the phase separation properties of the nucleolus in cells rely on continual activity of RNA Pol I, since its inhibition leads to disruption of these compartments(Ide et al. 2020). Our results point to a model in which *Dux* is held in repressive perinucleolar heterochromatin that is maintained through LLPS, with perturbation of rRNA transcription or direct inhibition of phase separation sufficient to cause *Dux* dissociation and de-repression. Similarly, we propose that the NPB structures of early 2C embryos and 2C-like cells are not competent for *Dux* repression. It will be interesting in future work to understand in further depth how the nucleolar periphery provides a repressive compartment like the nuclear lamina(Kind and van Steensel 2010). We previously found that LINE1 RNA and repressors Kap1 and Ncl bind both *Dux* and rDNA in ESCs(Percharde et al. 2018) with a recent report also identifying nucleolar Lin28 as a *Dux* repressor within this complex(Sun et al. 2021). Furthermore, it is likely that other repressor proteins may colocalise at the nucleolar periphery. For example, the histone methyltransferase, G9a has been reported in the nucleolus(Yuan et al. 2007), while a repressive role for nucleolar RNA Pol II itself has been recently discovered(Abraham et al. 2020).

In addition to *Dux* regulation, our work points to a wider role for nucleolar chromatin in gene regulation and its dynamic establishment in early embryos. RNA-seq data upon iPol I suggests that genes within Type II NADs, regions which are only associated with nucleoli and not constitutive LADs, are most sensitive to nucleolar disruption. In contrast, Type I NADs that show both nucleolar and constitutive lamina association are not upregulated upon acute iPol I, in agreement with their classification as constitutive heterochromatin and their low expression(Vertii et al. 2019; Bizhanova et al. 2020). Towards this, HDL treatment in MEFs was shown to cause relocalisation of a Type II, but not Type I, NAD(Vertii et al. 2019). Future work is needed to understand if these distinct NAD classes show differences in their association strength with the nucleolus, or if their activation depends on further mechanisms or factors upon dissociation. For example, neuronal NAD genes detach from the nucleolus upon neural progenitor cell differentiation yet do not yet become activated, supporting the model that their release might poise them for later expression(Bersaglieri et al. 2020).

More broadly, it will be important in future work to understand how nucleolar chromatin organisation proceeds in early embryos, as well as to uncover which genes rely on nucleolar association for their repression. Together with new findings of nucleolar function/dysfunction in multiple processes such as protein quality control(Azkanaz et al. 2019; Frottin et al. 2019), cancer(Lindstrom et al. 2018) and aging(Buchwalter and Hetzer 2017; Tiku and Antebi 2018), our data reveal a novel axis of nucleolar biology in early development and reflect the multi-faceted function of the nucleolus.

## Materials and Methods

### Mice and embryos

All animal experiments were performed according to a UK Home Office Project Licence in a Home Office-designated facility, using 4-6-week old female and 2-6-month-old male CD1 mice (Charles River). Animals were maintained on a 12h light/dark cycle and provided with *ad libitum* food and water in individually ventilated cages. Female mice were superovulated by intra-peritoneal injection of 5 I.U of pregnant mare serum gonadotropin (PMSG, Folligon, MSD Animal Health), followed by 5 I.U of human chorionic gonadotropin (hCG, Chorulon, MSD Animal Health) 46-48h later, then placed immediately with males. Zygotes were collected from oviducts ~22-24h post-hCG in M-2 medium (Sigma, M7167), isolated from cumulus cells with 200μg/mL Hyalurionidase (Sigma, H3506), washed through successive drops of M-2, and then cultured in pre-equilibrated KSOMaa (Sigma, MR-106-D) in microdrops overlaid with mineral oil (Sigma, M5310) or in 4-well dishes. Zygotes were cultured in a humidified incubator at 37oC, 5% CO2, until early 2-cell (31-33h post-hCG), late 2-cell (48-49h post-hCG), morula (3d post-hCG), or blastocyst (4d post-hCG).

### ESC culture

Mouse E14Tg2A (E14) ESCs (male) were used for all experiments(Hooper et al. 1987) and to derive 2C-GFP reporter cells. 2C-GFP reporter ESCs are described previously(Percharde et al. 2018) and were used when prior purification of larger numbers of 2C-like cells was not needed or for validation. All ESCs were cultured at 37oC, 5% CO2, on 0.1% gelatin-coated plates in ES-FBS culture medium (high glucose DMEM GlutaMAX with sodium pyruvate (Thermo Fisher Scientific), 15% FBS (Gibco), 0.1mM non- essential amino acids (Gibco), 0.1mM 2-Mercaptoethanol (Millipore) and 1,000U/ml LIF supplement (ESGRO, Millipore). ESCs were routinely tested for mycoplasma and found to be negative. Inhibitors (Table S2) were added to ESCs at the indicated concentrations unless otherwise explicitly mentioned in the figure legend.

### 2C-GFP/CD4 cell line

The 2C-GFP reporter construct(Ishiuchi et al. 2015) was modified to insert a T2A cleavage element followed by the extracellular portion of mouse CD4 (a.a.1-427) immediately downstream of GFP, so that activation of MERVL in the 2C-like state labels cells doubly positive for GFP and CD4. ESCs are negative for CD4 expression, enabling rapid purification of endogenous 2C-like cells via selection for CD4+ surface expression. E14 ESCs were nucleofected with 4ug linearized 2C-GFP plasmid and plated at low density in 10cm2 plates then selected with 250 μg/mLG418 (Mirus) for 8 days. Individual colonies were picked and expanded, with a single colony that showed high specific expression of GFP expanded and used for subsequent validations and experiments. For 2C-GFP/CD4 isolation, cells were trypsinised, washed, and resuspended in FACS buffer (PBS, 3% FBS, 1mM EDTA) then either isolated by MACS, using CD4 (L3T4) microbeads (Miltenyi Biotec), or with the EasySep mouse CD4 Positive selection kit II (StemCell), according to the manufacturer’s protocols in each case. Apart from Figure S1, all purification experiments were performed with the EasySep kit. Flow-through cells were collected as the 2C-negative population. For flow cytometry analysis, ESCs were pelleted and resuspended in FACS buffer containing 1:8000 Sytox Blue (Thermo Fisher Scientific) to enable exclusion of dead cells.

### siRNA-mediated knockdown

The nucleolar miniscreen was performed with a Cherry Pick custom library plate of OnTargetPlus siRNAs (Horizon Discovery), consisting of 20 wells of different gene-targeting siRNA pools and 3 siControl wells. 2C-GFP ESCs (non-CD4) at a density of 10,000 cells per 96-well were transfected in suspension with 3pmol siRNA and 0.17uL Lipofectamine 2000 per well of a 96-well plate and incubated overnight. The media was changed the following day then cells cultured for a further 2 days before analysis. For flow cytometry, ESCs were trypsinised, transferred to a 96-well, round-bottom plate, pelleted, washed and then resuspended in PBS plus 1:8000 Sytox Blue (Thermo Fisher Scientific), then the %GFP in live cells analysed by flow cytometry on a BD Fortessa cytometer. The nucleolar miniscreen was performed in triplicate wells and the entire experiment repeated on a different day with highly similar results. Z-scores were calculated as the %2C-GFP value for each factor minus the average 2C-GFP level for the entire plate, divided by the plate standard deviation. Other siRNA transfections or validation experiments were performed as above, scaling up cell numbers, Lipofectamine and siRNA amounts accordingly for ESCs cultured in 12- or 24-well plates, with cells harvested at the indicated time points.

### Nascent transcription/translation assays

Nascent transcription (EU) and translation (HPG) assays were carried out as described previously(Percharde et al. 2018) using Click-iT Assay Kits (Thermo Fisher Scientific) and according to the manufacturer’s protocol. For HPG assays, ESCs were cultured in medium made with DMEM lacking methionine (Gibco, #21013024) for 1h prior to HPG addition. ESCs were cultured with 1 mM EU or 50 uM HPG for 45 min before fixation, permeabilization and Click-iT reaction. Where indicated, immunofluorescence labelling was carried out prior to Click-iT as described above with the exception that primary antibodies were added for 1-2h at RT.

### Western blotting

Whole cell extracts were prepared from ESCs by scraping in ice-cold RIPA buffer containing protease-inhibitors (Halt), incubating for 30 min at 4oC, then pelleting at 16000g, 20 min to remove insoluble material. Proteins were quantified by the BCA assay (Pierce) and equal amounts loaded onto 4-12% Bolt Bis-Tris plus SDS-PAGE gels (Thermo Fisher) to separate proteins, then transferred onto PVDF membranes. Blocking was performed in 5% milk/PBS-T for 1h then membranes incubated overnight with primary antibodies at 4oC in milk/PBS-T. Next day, membranes were incubated with the appropriate HRP-conjugated secondary antibodies (Cell Signaling) for 1h, then proteins detected by ECL reagent on an Amersham Imager 680.

### RNA Extraction and Expression Analysis

RNA was isolated directly from ESCs by scraping in RLT lysis buffer (Qiagen) containing 1:100 beta-mercaptoethanol (Sigma), or RLT was added to equal numbers of ESCs for CNN approaches. RNA was purified and DNAse I treated according to the manufacturer’s instructions using RNeasy mini kits (Qiagen). For embryo inhibitor experiments, 2-cell embryos were flushed at 46h post-hCG and cultured in KSOMaa medium containing either 1μM CX-5461, BMH-21, or 0.1% DMSO in a 4-well dish. Culture in inhibitors began after 1h for a period of 8 or 24 hours. Equal numbers of embryos per experimental condition were lysed in 75 μL buffer RLT prepared as above, and the RNA isolated according to RNeasy micro kits (Qiagen). In ESCs and embryos, cDNA synthesis was performed with up to 1μg DNase-treated RNA using a High Capacity RNA-to-cDNA kit (Thermo Fisher Scientific), and qRT-PCR performed with SYBR green (KAPA) on a QuantStudio 5 qPCR machine (Thermo Fisher Scientific). qRT-PCR data were normalised to two housekeeping genes (Rpl7/H2A), unless a cell-number-normalisation (CNN) approach was used as detailed in the legend. Primer sequences are described in Table S2.

### RNA-sequencing and Analysis

For RNA-seq, RNA was extracted utilizing the RNeasy mini kit as for qRT-PCR, then 3 biological replicates per condition of DNase-treated total RNA spiked with ERCCs (Thermo Fisher) was used for RNA-seq library preparation and sequencing at the MRC LMS genomics core facility. RNA quality was assessed using the Agilent 2100 RNA 6000 Nano assay and libraries were prepared using the NEBNext Ultra II Directional RNA Library Prep Kit with NEBNext Poly(A) mRNA Magnetic Isolation Module, following manufacturer’s instructions. Library quality was evaluated using the Agilent 2100 High-Sensitivity DNA assay, and their concentrations measured using the Qubit™ dsDNA HS Assay Kit. Libraries were pooled in equimolar quantities and sequenced on an Illumina NextSeq 2000 to generate a minimum of 40 million Single Read 50bp reads (with unique 8bp dual indexes) per sample. Reads were trimmed and aligned to reference genome mm10 plus ERCCs using Tophat2. Default settings were used apart from the specification ‘g −1’ to map each multimapping read to one random TE or gene in the genome. Reads were counted using the Subread package, FeatureCounts to each gene or TE family. Data were filtered to exclude rows with counts per million (cpm) >0 in fewer than 3 samples. To account for any global decreases in RNA amounts due to iPol I, we used our previously described cell-number-normalised (CNN) approach to normalise reads to the abundance of ERCC spikeins(Percharde et al. 2017) using Limma Voom. All other RNA-seq analyses and statistics were performed in R/Bioconductor. Normalised RNA-seq expression data are available in Table S1. RNA-seq data have been uploaded to GEO, accession GSE185424.

For analysis of NAD gene expression, NAD regions defined in ESCs were taken from Bizhanova et al., 2020(Bizhanova et al. 2020). Type I NADs (overlapping constitutive LADs) and Type II NADs (overlapping constitutive interLADs) were defined as described in Vertii et al., 2019(Vertii et al. 2019) using LAD data from Peric-Hupkes et al., 2010(Peric-Hupkes et al. 2010), after LAD coordinates were shifted to mm10 using LiftOver tool (UCSC Genome Browser). For an example of NAD/LAD classification at the *Dux* locus see Figure S7. Ranked lists of log2 fold-change were prepared from RNA-seq of Pol I inhibitor treated ESCs (this study) and Ncl and LINE1 knockdown ESCs(Percharde et al. 2018). Ranked lists and Type I and Type II NAD files were submitted to the GSEA pre-ranked tool (genepattern.org) with the following parameters: permutations = 10000, collapse dataset = No_Collapse, and max gene set size = 4000. Normalised enrichment score (NES) and false discovery rate q-value (FDR) for each RNA-seq dataset were plotted using R/ggplot2.

### ESC Immunofluorescence

ESCs were allowed to adhere to Matrigel-coated 8-well chambers or 10 mm glass coverslips for 1-2h, fixed in 4% PFA for 10min, then stored in PBS until staining. Blocking and permeabilization was carried out in one step in immunofluorescence (IF) buffer (PBS, 10% donkey serum, 2.5% BSA), plus 0.4% Triton X-100 for 30min. Primary antibody incubations were carried out overnight at 4°C using the indicated antibodies and dilutions in IF buffer in Table S2. Next day, samples were washed with PBS and incubated in secondary antibodies (1:500 Alexa-488nm, 594nm, or 647nm-conjugated antibodies) for 1h at RT, followed by a wash for 30 min in PBS plus DAPI, two more washes in PBS, then samples mounted in Vectashield mounting medium containing DAPI. Confocal images were taken on a Leica SP5 fluorescent microscope under an oil immersion 63X objective.

### Embryo in vitro culture EU/HPG and IF experiments

Embryos were fixed in 4% PFA in PBS containing 0.1% Triton X-100 for 30 min, followed by three washes in PBS containing 0.1% PVA (PBS-PVA). Samples were permeabilised in PBS containing 0.5% Triton X-100 for 30 minutes, followed by blocking in 5% BSA in PBS for 1.5 hours. A 1:100 dilution of primary antibody (rabbit anti-nucleolin, Abcam, ab22558) was prepared in blocking solution, and embryos were incubated in 10 μl drops in a humidified Terasaki plate (Greiner Bio-One) at 4°C overnight. Embryos were washed three times in PBS containing 0.1% Tween-20 and 0.1% PVA (PBST-PVA) for five minutes. A 1:500 dilution of secondary antibody (donkey anti-rabbit Alexa Fluor 488, ThermoFisher, A21206) was prepared in blocking solution, and embryos were incubated in 10 μL drops in a humidified Terasaki plate for 1h in the dark. EU and HPG assays were performed as for ESCs, except that embryos were incubated for 1h in pre-equilibrated KSOM (without amino acids; Millipore), prior to incubation in KSOM containing 500 μm HPG or 1 mM EU for 2h. Embryos were fixed in 4% PFA in PBS containing 0.1% Triton X-100 for 15 minutes, and permeabilised in 0.5% Triton X-100 in PBS for 20 minutes, prior to Click-iT reactions. Prior to mounting, all samples were washed three times in PBST-PVA for five minutes, followed by a 30 min incubation in 1:1000 DAPI in PBS, and a further three washes in PBS-PVA. Embryos were mounted in Vectashield (Vector Laboratories) under a 20×20mm #1.5 coverslip (Agar Scientific) supported at the corners by Dow Corning high-vacuum silicone grease (Sigma-Aldrich) and sealed with nail polish. Confocal images were captured using a Leica SP5 or SP8 fluorescence microscope using an oil immersion 40X objective and acquired in 1 μm Z-stacks. All steps were carried out in 500 μl volumes at room temperature unless otherwise noted.

### Single Molecule RNA FISH combined with immunofluorescence staining (Immuno-smRNA-FISH)

*Dux* smRNA-FISH was based on branched DNA technology (*https://www.thermofisher.com/uk/en/home/life-science/cell-analysis/cellular-imaging/in-situ-hybridization-ish/rna-fish.html) to* detect the dynamics of *Dux* expression upon the inhibition of ribosomal RNA synthesis. A target set of 20 short Alexa647 labelled oligo mouse *Dux* ViewRNA ISH probe (VB6-3223670-VC, type-6; ThermoFisher Scientific) was designed and used to hybridize to *Dux* RNA sequence. Cells were first adhered on Matrigel-coated 10 mm glass coverslips and fixed with 4% paraformaldehyde in DEPC-treated PBS, containing 2.5% Acetic acid for 10 mins. Immunolabeling with performed with mouse anti-B23 primary antibody (1:300; 2h) then detected with Alexa 488 donkey anti-mouse antibody (Invitrogen; 1:500; 1h), then further fixed with 4% paraformaldehyde in DEPC-PBS, 10 min to preserve immunocomplexes before FISH. smRNA FISH was carried out in accordance with the manufacturer’s protocol for the ViewRNA TM ISH Cell Assay (QVC001; ThermoFisher Scientific). Briefly, previously immunolabled cells were washed in PBS (3x), permeabilised (1:2 digestion solution in DEPC-PBS; 5 min), protease treated (1:8000 in DEPC-PBS, 10 min), and subsequently hybridized with Alexa 647 ViewRNA ISH *Dux* probe (P4, 1:100 in QF diluent; 3h, 40oC). Following probe hybridisation, cells were washed and then subjected to sequential hybridisation with pre-amplifier DNA, amplifier DNA and fluorophore labeled in provided diluent (1:25, 1h, 40°C each step). After detection, cells were washed and nuclei were stained with 1μg/ml DAPI in PBS before imaging. Control experiments were performed with either hybridisation with mouse B-actin probe as a positive control that revealed abundant b-actin RNA foci throughout nucleus and cytoplasm, or Rnase A treatment (250 μg/ml in PBS, 2h, 37oC) prior to hybridisation, which abrograted *Dux* signals.

### DNA FISH with immunofluorescence (Immuno-DNA-FISH)

Immunofluorescence detection of nucleolus and nuclear lamina combined with DNA was performed essentially as described previously(Beagrie et al. 2017). Briefly, ESCs were fixed with 4% plus 0.1% Triton in PBS, 10 min, washed in PBS (3x), equilibrated in 20% glycerol in PBS (3x, 10min), subsequently then frozen in liquid nitrogen and stored at −80°C. After thawing, cells were washed in PBS 3x, permeabilized for 10 min with 0.1% triton in PBS and blocked with 2% BSA-PBS, 30 min, before immunolabeling. Cells were incubated with the indicated mouse anti-B23 as above; and rabbit anti-laminB1 (1:2000; ab16048; Abcam) antibodies in 2% BSA in PBS for 2h, then detected with AlexaFluor488 or 647-conjugated antibodies. After immunolabelling, cells were fixed with 4% PFA in PBS for 30 min prior to FISH to preserve immunocomplexes during FISH. *Dux* oligo probes used to detect *Dux* foci were used and labelled with Cy3 fluorphores (PA23001; Amersham) as described previously (Percharde et al. 2018). For hybridisation, 1 μg mouse Cot1 DNA (18440; Invitrogen), 10 μg salmon sperm DNA (15632011; Invitrogen) and 3 μl Cy3-labelled Dux oligos, respectively, were precipitated and resuspended in 6 μl of hybridisation buffer (H7782, Sigma-Aldrich) ready for DNA FISH. Immunolablled cells were rinsed 3x in PBS, incubated for 15 min in 20 mM glycine in PBS, rinsed 3x in PBS, permeabilised for 10 min with 0.2% Triton, and then washed again. Cells were incubated for 1-2h at 37°C with 250 μg/ml RNase A (Sigma) in 2x SSC, treated for 10 min with 0.1 M HCl, dehydrated in ethanol (50% to 100% series, 3 min each), dried briefly, denatured for 10 min at 80°C in 70% deionized formamide in 2xSSC, and then re-dehydrated as above. After a brief period of drying, coverslips were overlaid onto probe mixture on Hybrislips (H18200; Molecular Probe by Life Technology) and sealed with Fixogum rubber cement (11FIXO0125; MP Biomedicals) for in situ hybridisation. Hybridization was carried out at 37°C in a moist chamber for at least 40 h. Post-hybridization washes were as follows: 40% formamide in 2xSSC (3x 10min); 2xSSC (3x, 10min); and (3x, PBS). Nuclei were counterstained with 1μg/ml DAPI in PBS for 30 min and mounted in (Vectashield H-1000; Vector Laboratories) immediately prior to imaging.

### Super-Resolution Structured Illumination Microscopy (SIM)

Purified 2C-GFP/CD4-negative and positive cells or wild-type ESCs incubated for 8h with or without iPol I were adhered on Matrigel-coated μ-Slide 8 Well Glass Bottom (80827; Ibidi) and fixed with 4% paraformaldehyde in PBS for 10 min then stored in PBS at 4°C before immunolabeling. Cells were rinsed 3x in PBS, incubated for 15 min in 20 mM glycine in PBS, rinsed, permeabilized with 0.1% Triton X-100 for 10 min in PBS, blocked for 1h with 4% BSA in PBS then incubated with mouse anti-B23 to detect nucleoli in 4% BSA in PBS overnight at 4oC. Next day, samples were washed 3x for 60 min with 2% BSA in PBS, then incubated 2h with Alexa secondary antibodies (1:250) washed again as above. Finally, samples were washed and counterstained with 1μg/ml DAPI (30 min), rinsed successively in PBS before coverslips were mounted in VectaShield. The long incubation times used allow for antibody accessibility throughout the cells, providing the highest sensitivity for SIM imaging. Multi-colour SIM imaging was performed using a Zeiss Elyra S.1 (Carl Zeiss Microimaging) and a Plan-Apochromat 63×/1.4 oil lens. Raw SIM images were acquired with an sCMOS camera (pco.Edge 4.2) using five phase shifts and three grid rotations, with a z step size of 0.1 μm. Different fluorescent labels were acquired sequentially using 642, 561, 488 and 405nm laser lines. SIM images were reconstructed with ZEN 2012 SP4 (Black) software (Carl Zeiss Microimaging, version 13.0.2.518), using default parameter settings. Channel alignment was performed using calibrations obtained from a multi-coloured bead slide, acquired with equivalent acquisition settings.

### ESC Confocal microscopy and Quantitative Image Analysis

Multi-colour image single snapshots (IF and EU/HPG assays) or Z-stacks (250nm stepsize, RNA/DNA FISH) were acquired with a laser-scanning confocal microscope with a pinhole diameter of 1 Airy unit (Leica TCS SP5 or SP8; Objective lens: 63x/1.40NA Oil CS2 HC PL APO; laser lines: 405/488/552/638nm). Different channels were imaged sequentially to avoid bleed through and cross-excitation and then exported as TIFF files for further images analysis. RNA and DNA FISH image Z-stacks were also acquired on an Olympus spinning disk confocal system based on an IX83 inverted microscope stand (Yokogawa CSU-W1 scanhead with 50μm diameter pinhole disk; Objective lens: 60x/1.40NA Plan-Apo; Hamamatsu ORCA Flash 4.0 V2 camera, stepsize 200nm). Raw .lif images were processed into TIFF files and merged, each channel manually thresholding or filtering with the same setting in Fiji software for data analysis.

Dux FISH 3D spatial analysis was undertaken using custom written scripts in Fiji for nuclei and nucleoli segmentation, and Imaris (version 9.6.0, Bitplane AG) for Dux FISH probe identification and distance measurements. Briefly, identification of LaminB1 labelled Nuclei and B23 labelled Nucleoli was performed in Fiji in combination with the MorphoLibJ plugin to perform 3D segmentation. Labelled image masks generated were combined with the original image stack and imported into Imaris for use with the ‘Cells’ Package, to facilitate interactive review of 3D segmentation results and Dux FISH probe identification. Identified *Dux* FISH loci were related to each individual nucleus and contained nucleoli, and distance to the nearest nucleoli and nuclei border measured.

The analysis of nucleolar morphology and fluorescence intensities was carried out with a custom written CellProfiler pipeline (https://cellprofiler.org/; software version 4.1.3). To describe and quantify the number of “ring-shaped” nucleoli - showing a distinct fluorescent B23 signal at the rim of the nucleoli with a much dimmer interior signal – the fluorescence intensity distribution over four concentric layers in each nucleolus was measured. The mean fluorescence intensity of the outer layer was divided by the mean intensity of the innermost layer and any nucleolus with a ratio greater than 1.4 was counted in the “ring-shaped” category. Nucleolar circularity (“Form Factor”) was calculated within CellProfiler according to 4π × [Area]/[Perimeter]^2^, excluding very small nucleoli which are liable to give unreliable measurements (FormFactor values >1).

### Statistical analysis

All statistical analyses were carried out in Prism 7 or above (Graphpad) or R (RNA-seq data). Details of individual tests are outlined within each figure legend, including number and type of replication performed (n) and the reported error either as standard deviation (s.d) or standard error of the mean (s.e.m). All statistics are * p < 0.05, ** p < 0.01, *** p < 0.001, **** p < 0.0001 with the relevant test performed described in figure legends and corrections for multiple testing applied where necessary. Welch’s correction was applied to t-tests when the variance was unequal between conditions.

### Data Availability

RNA-seq data have been uploaded to GEO, accession GSE185424.

### Code Availability

RNA-seq data were analysed with standard packages and programs, as detailed in the Methods. Code for data processing and analysis are available at https://github.com/mpercharde/RNAseq and/or are available upon request.

## Acknowledgements

We thank Alexis Barr, Aydan Bulut-Karslioglu, Harry Leitch and members of the Percharde lab for critical reading and comments on the manuscript. We thank the MRC LMS genomics, imaging and flow cytometry cores for technical services. Thank you to Tristan Rodriguez for *p53*-null ESCs. Work in the Percharde lab is supported by a UKRI Future Leaders Fellowship (MC_EX_MR/S015930/1) and funding from the Medical Research Council (MRC, MC_UP_1605/4) to M.P. Work in the McManus Lab is supported by NIH 5R01GM123556.

## Author Contributions

M.P. conceived the project. S.Q.X., B.J.L, P.C, F.G-L, N.T-F.C and M.P. designed and performed experiments. C.W. and D.D. performed microscopy image analysis with S.Q.X,. R.T.W. generated 2C-GFP/CD4 ESCs, supervised by M.T.M., B.J.L and M.P. performed computational analysis. S.Q.X and M.P. wrote the manuscript with input from all authors.

## References

Abraham KJ, Khosraviani N, Chan JNY, Gorthi A, Samman A, Zhao DY, Wang M, Bokros M, Vidya E, Ostrowski LA et al. 2020. Nucleolar RNA polymerase II drives ribosome biogenesis. Nature 585: 298–302.

Azkanaz M, Rodriguez Lopez A, de Boer B, Huiting W, Angrand PO, Vellenga E, Kampinga HH, Bergink S, Martens JH, Schuringa JJ et al. 2019. Protein quality control in the nucleolus safeguards recovery of epigenetic regulators after heat shock. Elife 8.

Beagrie RA, Scialdone A, Schueler M, Kraemer DC, Chotalia M, Xie SQ, Barbieri M, de Santiago I, Lavitas LM, Branco MR et al. 2017. Complex multi-enhancer contacts captured by genome architecture mapping. Nature 543: 519–524.

Bersaglieri C, Kresoja-Rakic J, Gupta S, Bär D, Kuzyakiv R, Santoro R. 2020. Genome-wide maps of nucleolus interactions reveal distinct layers of repressive chromatin domains. bioRxiv: 2020.2011.2017.386797.

Binek A, Rojo D, Godzien J, Ruperez FJ, Nunez V, Jorge I, Ricote M, Vazquez J, Barbas C. 2019. Flow Cytometry Has a Significant Impact on the Cellular Metabolome. J Proteome Res 18: 169–181.

Bizhanova A, Yan A, Yu J, Zhu LJ, Kaufman PD. 2020. Distinct features of nucleolus-associated domains in mouse embryonic stem cells. Chromosoma 129: 121–139.

Borsos M, Torres-Padilla ME. 2016. Building up the nucleus: nuclear organization in the establishment of totipotency and pluripotency during mammalian development. Genes & development 30: 611–621.

Boskovic A, Eid A, Pontabry J, Ishiuchi T, Spiegelhalter C, Raghu Ram EV, Meshorer E, Torres-Padilla ME. 2014. Higher chromatin mobility supports totipotency and precedes pluripotency in vivo. Genes & development 28: 1042–1047.

Bosnakovski D, Gearhart MD, Ho Choi S, Kyba M. 2021. Dux facilitates post-implantation development, but is not essential for zygotic genome activationdagger. Biol Reprod 104: 83–93.

Buchwalter A, Hetzer MW. 2017. Nucleolar expansion and elevated protein translation in premature aging. Nat Commun 8: 328.

Bywater MJ, Poortinga G, Sanij E, Hein N, Peck A, Cullinane C, Wall M, Cluse L, Drygin D, Anderes K et al. 2012. Inhibition of RNA polymerase I as a therapeutic strategy to promote cancer-specific activation of p53. Cancer Cell 22: 51–65.

Casser E, Israel S, Witten A, Schulte K, Schlatt S, Nordhoff V, Boiani M. 2017. Totipotency segregates between the sister blastomeres of two-cell stage mouse embryos. Sci Rep 7: 8299.

Chen Z, Zhang Y. 2019. Loss of DUX causes minor defects in zygotic genome activation and is compatible with mouse development. Nature genetics 51: 947–951.

Choi YJ, Lin CP, Risso D, Chen S, Kim TA, Tan MH, Li JB, Wu Y, Chen C, Xuan Z et al. 2017. Deficiency of microRNA miR-34a expands cell fate potential in pluripotent stem cells. Science 355.

Chuong EB, Rumi MA, Soares MJ, Baker JC. 2013. Endogenous retroviruses function as species-specific enhancer elements in the placenta. Nature genetics 45: 325–329.

De Iaco A, Planet E, Coluccio A, Verp S, Duc J, Trono D. 2017. DUX-family transcription factors regulate zygotic genome activation in placental mammals. Nature genetics 49: 941–945.

De Iaco A, Verp S, Offner S, Grun D, Trono D. 2020. DUX is a non-essential synchronizer of zygotic genome activation. Development 147.

Dixit M, Ansseau E, Tassin A, Winokur S, Shi R, Qian H, Sauvage S, Matteotti C, van Acker AM, Leo O et al. 2007. DUX4, a candidate gene of facioscapulohumeral muscular dystrophy, encodes a transcriptional activator of PITX1. Proc Natl Acad Sci U S A 104: 18157–18162.

Eckersley-Maslin M, Alda-Catalinas C, Blotenburg M, Kreibich E, Krueger C, Reik W. 2019. Dppa2 and Dppa4 directly regulate the Dux-driven zygotic transcriptional program. Genes & development 33: 194–208.

Falahati H, Pelham-Webb B, Blythe S, Wieschaus E. 2016. Nucleation by rRNA Dictates the Precision of Nucleolus Assembly. Curr Biol 26: 277–285.

Fedoriw AM, Starmer J, Yee D, Magnuson T. 2012. Nucleolar association and transcriptional inhibition through 5S rDNA in mammals. PLoS Genet 8: e1002468.

Feric M, Vaidya N, Harmon TS, Mitrea DM, Zhu L, Richardson TM, Kriwacki RW, Pappu RV, Brangwynne CP. 2016. Coexisting Liquid Phases Underlie Nucleolar Subcompartments. Cell 165: 1686–1697.

Flechon JE, Kopecny V. 1998. The nature of the ‘nucleolus precursor body’ in early preimplantation embryos: a review of fine-structure cytochemical, immunocytochemical and autoradiographic data related to nucleolar function. Zygote 6: 183–191.

Frottin F, Schueder F, Tiwary S, Gupta R, Korner R, Schlichthaerle T, Cox J, Jungmann R, Hartl FU, Hipp MS. 2019. The nucleolus functions as a phase-separated protein quality control compartment. Science 365: 342–347.

Fulka H, Langerova A. 2014. The maternal nucleolus plays a key role in centromere satellite maintenance during the oocyte to embryo transition. Development 141: 1694–1704.

Gabellini D, Green MR, Tupler R. 2002. Inappropriate gene activation in FSHD: a repressor complex binds a chromosomal repeat deleted in dystrophic muscle. Cell 110: 339–348.

Geng LN, Yao Z, Snider L, Fong AP, Cech JN, Young JM, van der Maarel SM, Ruzzo WL, Gentleman RC, Tawil R et al. 2012. DUX4 activates germline genes, retroelements, and immune mediators: implications for facioscapulohumeral dystrophy. Dev Cell 22: 38–51.

Ginisty H, Amalric F, Bouvet P. 1998. Nucleolin functions in the first step of ribosomal RNA processing. EMBO J 17: 1476–1486.

Grow EJ, Weaver BD, Smith CM, Guo J, Stein P, Shadle SC, Hendrickson PG, Johnson NE, Butterfield RJ, Menafra R et al. 2021. p53 convergently activates Dux/DUX4 in embryonic stem cells and in facioscapulohumeral muscular dystrophy cell models. Nature genetics 53: 1207–1220.

Guallar D, Bi X, Pardavila JA, Huang X, Saenz C, Shi X, Zhou H, Faiola F, Ding J, Haruehanroengra P et al. 2018. RNA-dependent chromatin targeting of TET2 for endogenous retrovirus control in pluripotent stem cells. Nature genetics 50: 443–451.

Guo M, Zhang Y, Zhou J, Bi Y, Xu J, Xu C, Kou X, Zhao Y, Li Y, Tu Z et al. 2019. Precise temporal regulation of Dux is important for embryo development. Cell Res 29: 956–959.

Haddach M, Schwaebe MK, Michaux J, Nagasawa J, O’Brien SE, Whitten JP, Pierre F, Kerdoncuff P, Darjania L, Stansfield R et al. 2012. Discovery of CX-5461, the First Direct and Selective Inhibitor of RNA Polymerase I, for Cancer Therapeutics. ACS Med Chem Lett 3: 602–606.

Hendrickson PG, Dorais JA, Grow EJ, Whiddon JL, Lim JW, Wike CL, Weaver BD, Pflueger C, Emery BR, Wilcox AL et al. 2017. Conserved roles of mouse DUX and human DUX4 in activating cleavage-stage genes and MERVL/HERVL retrotransposons. Nature genetics 49: 925–934.

Hooper M, Hardy K, Handyside A, Hunter S, Monk M. 1987. HPRT-deficient (Lesch-Nyhan) mouse embryos derived from germline colonization by cultured cells. Nature 326: 292–295.

Hu Z, Tan DEK, Chia G, Tan H, Leong HF, Chen BJ, Lau MS, Tan KYS, Bi X, Yang D et al. 2020. Maternal factor NELFA drives a 2C-like state in mouse embryonic stem cells. Nat Cell Biol 22: 175–186.

Huang N, Negi S, Szebeni A, Olson MO. 2005. Protein NPM3 interacts with the multifunctional nucleolar protein B23/nucleophosmin and inhibits ribosome biogenesis. J Biol Chem 280: 5496–5502.

Huang Y, Kim JK, Do DV, Lee C, Penfold CA, Zylicz JJ, Marioni JC, Hackett JA, Surani MA. 2017. STELLA modulates transcriptional and endogenous retrovirus programs during maternal-to-zygotic transition. Elife 6.

Ide S, Imai R, Ochi H, Maeshima K. 2020. Transcriptional suppression of ribosomal DNA with phase separation. Sci Adv 6.

Ishiuchi T, Enriquez-Gasca R, Mizutani E, Boskovic A, Ziegler-Birling C, Rodriguez-Terrones D, Wakayama T, Vaquerizas JM, Torres-Padilla ME. 2015. Early embryonic-like cells are induced by downregulating replication-dependent chromatin assembly. Nat Struct Mol Biol 22: 662–671.

James A, Wang Y, Raje H, Rosby R, DiMario P. 2014. Nucleolar stress with and without p53. Nucleus 5: 402–426.

Kind J, van Steensel B. 2010. Genome-nuclear lamina interactions and gene regulation. Curr Opin Cell Biol 22: 320–325.

Kunarso G, Chia NY, Jeyakani J, Hwang C, Lu X, Chan YS, Ng HH, Bourque G. 2010. Transposable elements have rewired the core regulatory network of human embryonic stem cells. Nature genetics 42: 631–634.

Kyogoku H, Kitajima TS, Miyano T. 2014. Nucleolus precursor body (NPB): a distinct structure in mammalian oocytes and zygotes. Nucleus 5: 493–498.

Laferte A, Favry E, Sentenac A, Riva M, Carles C, Chedin S. 2006. The transcriptional activity of RNA polymerase I is a key determinant for the level of all ribosome components. Genes & development 20: 2030–2040.

Lindstrom MS, Jurada D, Bursac S, Orsolic I, Bartek J, Volarevic S. 2018. Nucleolus as an emerging hub in maintenance of genome stability and cancer pathogenesis. Oncogene 37: 2351–2366.

Lu JY, Shao W, Chang L, Yin Y, Li T, Zhang H, Hong Y, Percharde M, Guo L, Wu Z et al. 2020. Genomic Repeats Categorize Genes with Distinct Functions for Orchestrated Regulation. Cell Rep 30: 3296–3311 e3295.

Macfarlan TS, Gifford WD, Agarwal S, Driscoll S, Lettieri K, Wang J, Andrews SE, Franco L, Rosenfeld MG, Ren B et al. 2011. Endogenous retroviruses and neighboring genes are coordinately repressed by LSD1/KDM1A. Genes & development 25: 594–607.

Macfarlan TS, Gifford WD, Driscoll S, Lettieri K, Rowe HM, Bonanomi D, Firth A, Singer O, Trono D, Pfaff SL. 2012. Embryonic stem cell potency fluctuates with endogenous retrovirus activity. Nature 487: 57–63.

Martin C, Brochard V, Migne C, Zink D, Debey P, Beaujean N. 2006. Architectural reorganization of the nuclei upon transfer into oocytes accompanies genome reprogramming. Mol Reprod Dev 73: 1102–1111.

Martinez Arias A, Nichols J, Schroter C. 2013. A molecular basis for developmental plasticity in early mammalian embryos. Development 140: 3499–3510.

Nemeth A, Conesa A, Santoyo-Lopez J, Medina I, Montaner D, Peterfia B, Solovei I, Cremer T, Dopazo J, Langst G. 2010. Initial genomics of the human nucleolus. PLoS Genet 6: e1000889.

Ogushi S, Palmieri C, Fulka H, Saitou M, Miyano T, Fulka J, Jr. 2008. The maternal nucleolus is essential for early embryonic development in mammals. Science 319: 613–616.

Olbrich T, Vega-Sendino M, Tillo D, Wu W, Zolnerowich N, Pavani R, Tran AD, Domingo CN, Franco M, Markiewicz-Potoczny M et al. 2021. CTCF is a barrier for 2C-like reprogramming. Nat Commun 12: 4856.

Peaston AE, Evsikov AV, Graber JH, de Vries WN, Holbrook AE, Solter D, Knowles BB. 2004. Retrotransposons regulate host genes in mouse oocytes and preimplantation embryos. Dev Cell 7: 597–606.

Peltonen K, Colis L, Liu H, Trivedi R, Moubarek MS, Moore HM, Bai B, Rudek MA, Bieberich CJ, Laiho M. 2014. A targeting modality for destruction of RNA polymerase I that possesses anticancer activity. Cancer Cell 25: 77–90.

Percharde M, Lin CJ, Yin Y, Guan J, Peixoto GA, Bulut-Karslioglu A, Biechele S, Huang B, Shen X, Ramalho-Santos M. 2018. A LINE1-Nucleolin Partnership Regulates Early Development and ESC Identity. Cell 174: 391–405 e319.

Percharde M, Wong P, Ramalho-Santos M. 2017. Global Hypertranscription in the Mouse Embryonic Germline. Cell Rep 19: 1987–1996.

Peric-Hupkes D, Meuleman W, Pagie L, Bruggeman SW, Solovei I, Brugman W, Graf S, Flicek P, Kerkhoven RM, van Lohuizen M et al. 2010. Molecular maps of the reorganization of genome-nuclear lamina interactions during differentiation. Mol Cell 38: 603–613.

Quinodoz SA, Ollikainen N, Tabak B, Palla A, Schmidt JM, Detmar E, Lai MM, Shishkin AA, Bhat P, Takei Y et al. 2018. Higher-Order Inter-chromosomal Hubs Shape 3D Genome Organization in the Nucleus. Cell 174: 744–757 e724.

Ribbeck K, Gorlich D. 2002. The permeability barrier of nuclear pore complexes appears to operate via hydrophobic exclusion. EMBO J 21: 2664–2671.

Rossant J, Chazaud C, Yamanaka Y. 2003. Lineage allocation and asymmetries in the early mouse embryo. Philos Trans R Soc Lond B Biol Sci 358: 1341–1348; discussion 1349.

Rubbi CP, Milner J. 2003. Disruption of the nucleolus mediates stabilization of p53 in response to DNA damage and other stresses. EMBO J 22: 6068–6077.

Shadle SC, Zhong JW, Campbell AE, Conerly ML, Jagannathan S, Wong CJ, Morello TD, van der Maarel SM, Tapscott SJ. 2017. DUX4-induced dsRNA and MYC mRNA stabilization activate apoptotic pathways in human cell models of facioscapulohumeral dystrophy. PLoS Genet 13: e1006658.

Shav-Tal Y, Blechman J, Darzacq X, Montagna C, Dye BT, Patton JG, Singer RH, Zipori D. 2005. Dynamic sorting of nuclear components into distinct nucleolar caps during transcriptional inhibition. Mol Biol Cell 16: 2395–2413.

Sun Z, Yu H, Zhao J, Tan T, Pan H, Zhu Y, Chen L, Zhang C, Zhang L, Lei A et al. 2021. LIN28 coordinately promotes nucleolar/ribosomal functions and represses the 2C-like transcriptional program in pluripotent stem cells. Protein Cell.

Sundaram V, Cheng Y, Ma Z, Li D, Xing X, Edge P, Snyder MP, Wang T. 2014. Widespread contribution of transposable elements to the innovation of gene regulatory networks. Genome research 24: 1963–1976.

Svoboda P, Stein P, Anger M, Bernstein E, Hannon GJ, Schultz RM. 2004. RNAi and expression of retrotransposons MuERV-L and IAP in preimplantation mouse embryos. Dev Biol 269: 276–285.

Tarkowski AK. 1959. Experiments on the development of isolated blastomers of mouse eggs. Nature 184: 1286–1287.

Tiku V, Antebi A. 2018. Nucleolar Function in Lifespan Regulation. Trends Cell Biol 28: 662–672.

Vertii A, Ou J, Yu J, Yan A, Pages H, Liu H, Zhu LJ, Kaufman PD. 2019. Two contrasting classes of nucleolus-associated domains in mouse fibroblast heterochromatin. Genome research 29: 1235–1249.

Whiddon JL, Langford AT, Wong CJ, Zhong JW, Tapscott SJ. 2017. Conservation and innovation in the DUX4-family gene network. Nature genetics 49: 935–940.

Yang F, Huang X, Zang R, Chen J, Fidalgo M, Sanchez-Priego C, Yang J, Caichen A, Ma F, Macfarlan T et al. 2020. DUX-miR-344-ZMYM2-Mediated Activation of MERVL LTRs Induces a Totipotent 2C-like State. Cell Stem Cell 26: 234–250 e237.

Yao RW, Xu G, Wang Y, Shan L, Luan PF, Wang Y, Wu M, Yang LZ, Xing YH, Yang L et al. 2019. Nascent Pre-rRNA Sorting via Phase Separation Drives the Assembly of Dense Fibrillar Components in the Human Nucleolus. Mol Cell 76: 767–783 e711.

Yuan X, Feng W, Imhof A, Grummt I, Zhou Y. 2007. Activation of RNA polymerase I transcription by cockayne syndrome group B protein and histone methyltransferase G9a. Mol Cell 27: 585–595.

Zuccotti M, Garagna S, Merico V, Monti M, Alberto Redi C. 2005. Chromatin organisation and nuclear architecture in growing mouse oocytes. Mol Cell Endocrinol 234: 11–17.

