## Supplemental_data for "Nucleolar-based *Dux* repression is essential for 2-cell stage exit"

**Supplemental Information:**

Figures 1-7
Table S1: RNA-seq data in this study upon iPol I
Table S2: Reagents, probes and primer sequences used in this study

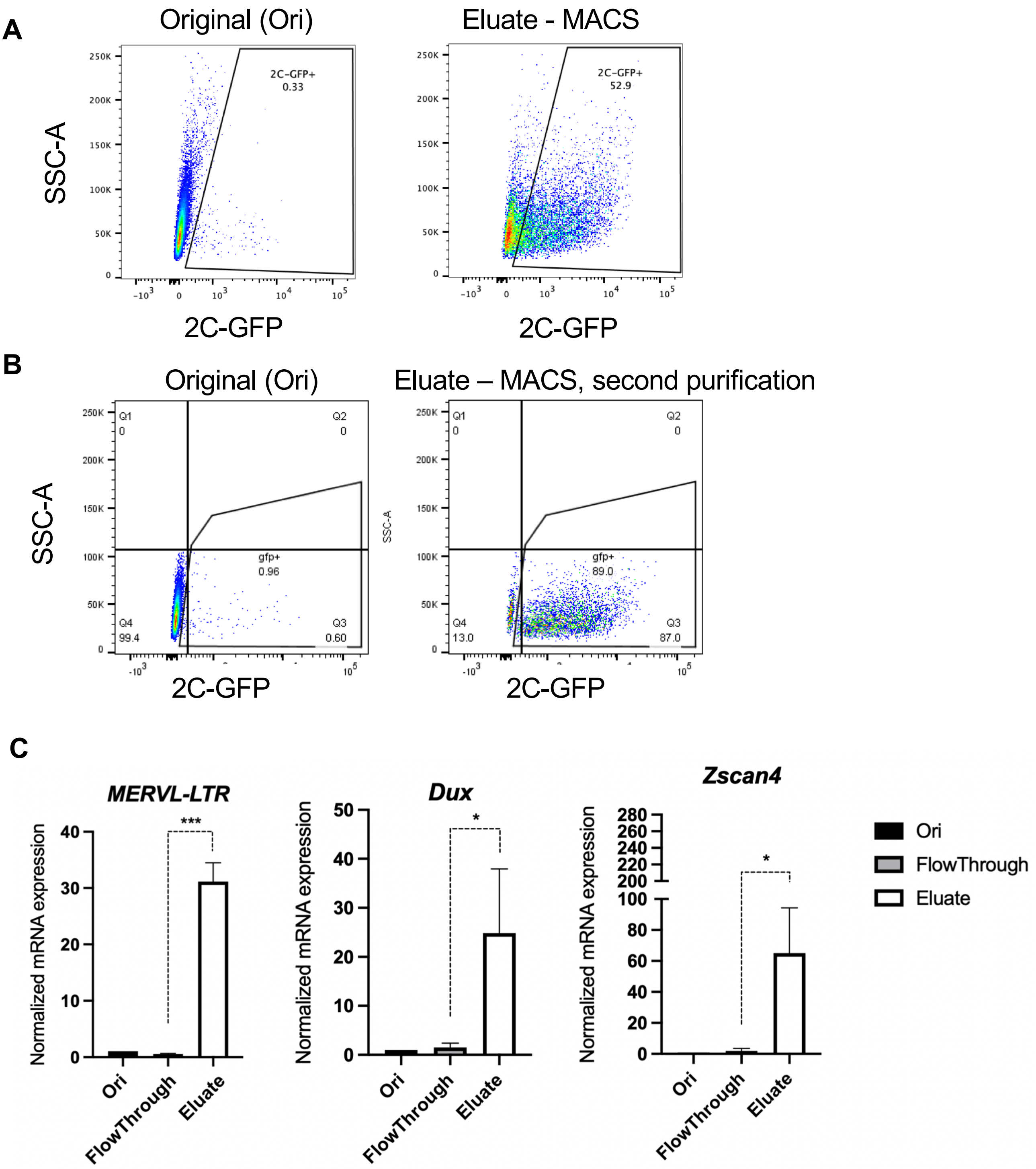


**Figure S1**. MACS purification of 2C-like cells.

A) Representative flow cytometry plots of 2C-GFP/CD4 ESCs before and after MACS purification

B) Flow cytometry plots showing increased purity of 2C-GFP+ cells after two rounds of MACS purification.

C) qRT-PCR analysis of 2C-specific transcripts after MACS purification. Note that 2C gene induction with MACS is lower than CD4 column-free, bead-based purification (StemCell Inc) (Figure 1D). CD4 column-free isolation was therefore used in all subsequent experiments throughout this study. Data are mean +/- s.e.m, 3 independent experiments, P values, unpaired Student's t-test.

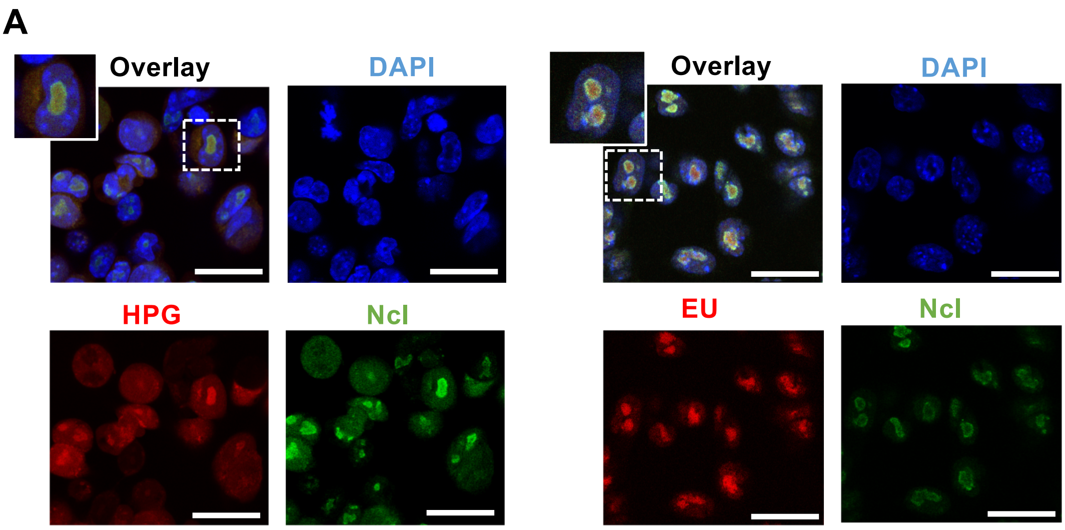


**Figure S2**. Nascent nucleolar RNA and protein expression.

A) Representative confocal microscopy images confirming overlap of Nucleolin (Ncl) and HPG (left) or EU (right) in ESCs. Scale, 25 µm.

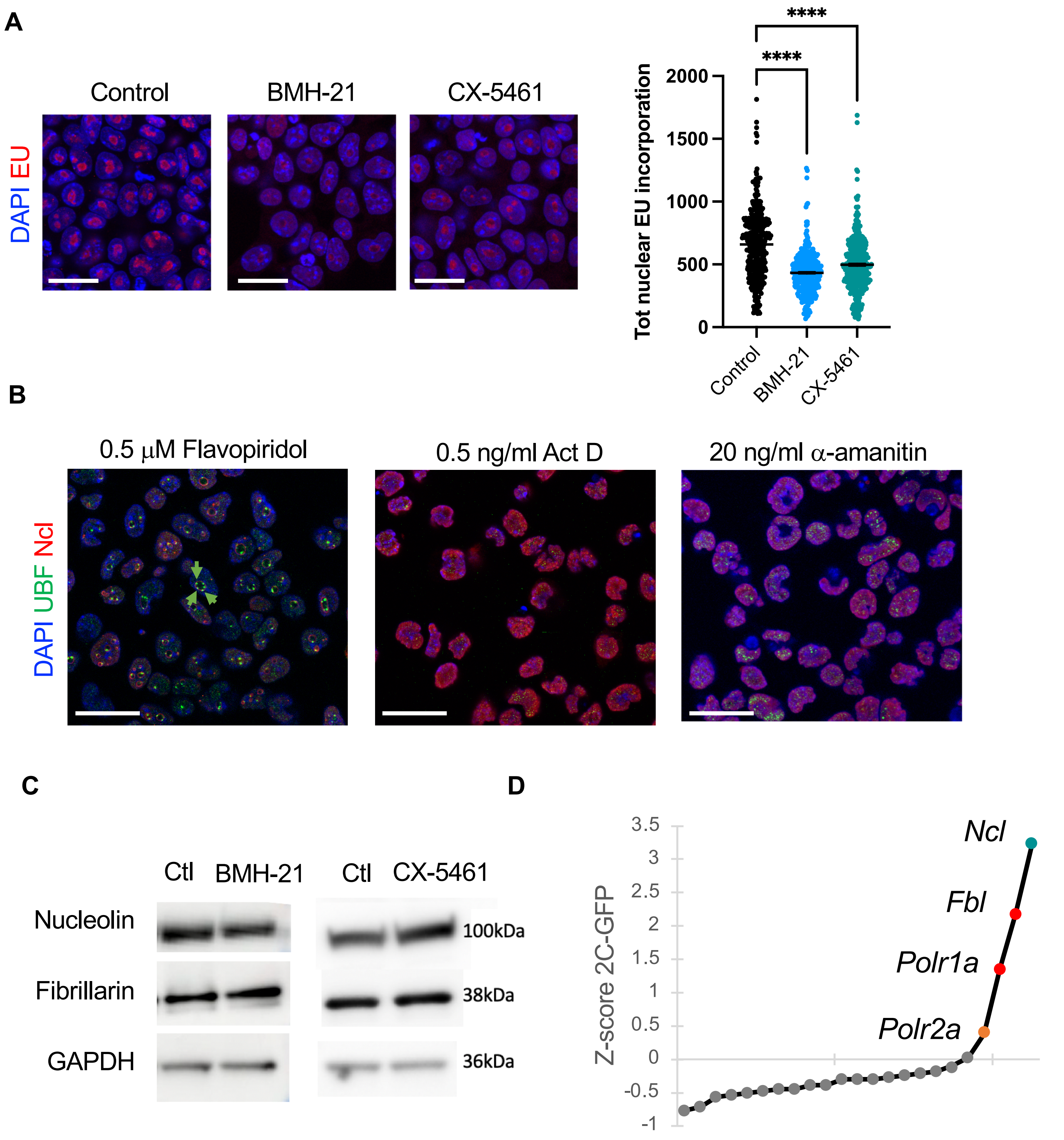


**Figure S3**. Nucleolar disruption by RNA Pol I inhibition

A) Representative confocal microscopy images and quantification of nascent RNA levels (EU incorporation) following 2h RNA Pol I inhibition. Data are mean +/- s.e.m, P values, one-way ANOVA with Dunnett multiple comparisons correction. Scale bars, 25 μm.

B) Representative confocal microscopy images of ESC nucleolar proteins and morphology 8h following the indicated Pol I/II inhibitors. Nucleolar caps are indicated with arrowheads. Scale bars, 25 μm.

C) Western blots of the indicated nucleolar proteins upon iPol I. Samples are representative of 2 independent experiments.

D) Z-scores for 2C-GFP cell induction following nucleolar protein knockdown, at d3 following transfection.

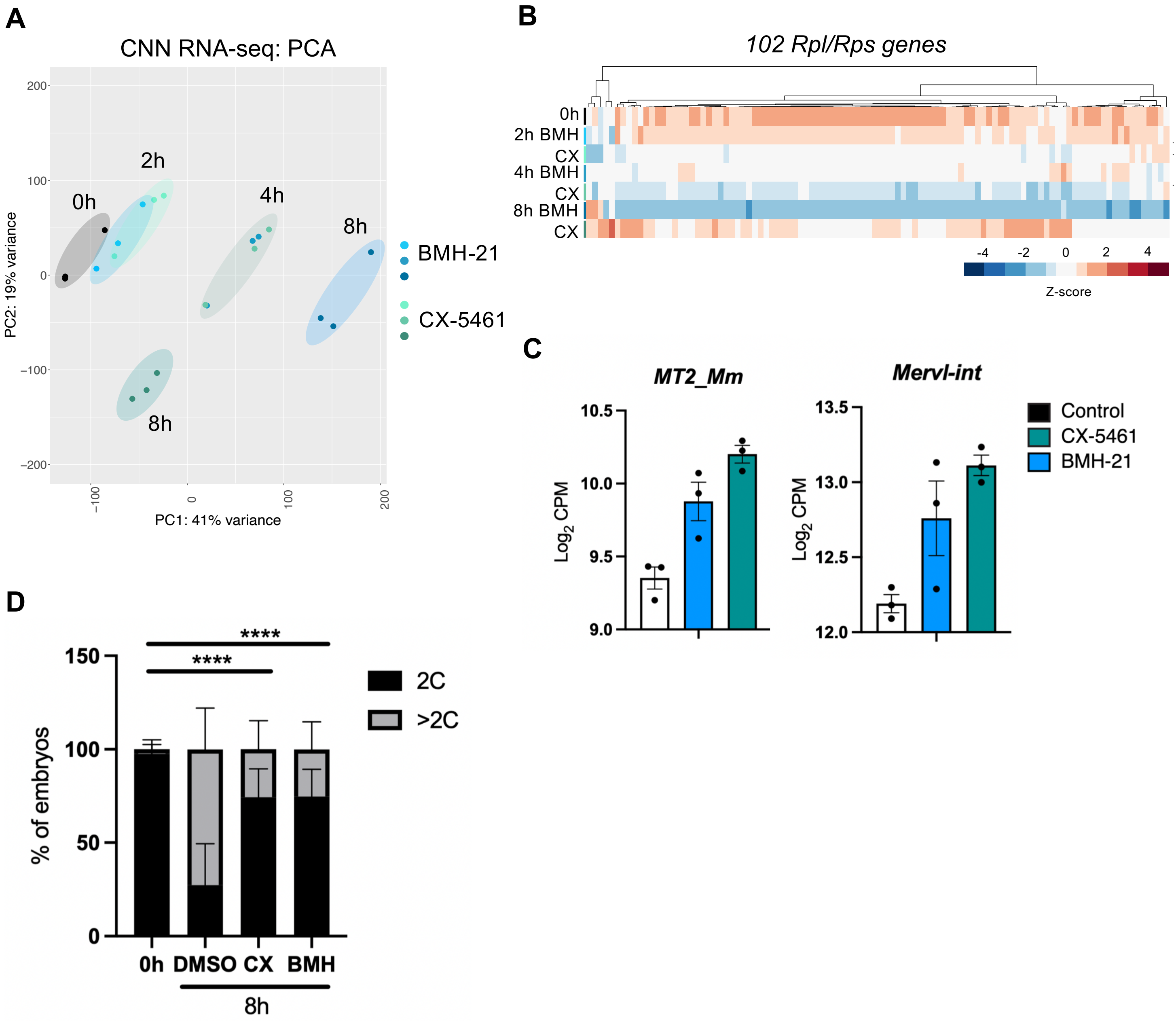


**Figure S4**. *Dux* activation upon iPol I

A) PCA of the first 2 principle components of CNN RNA-seq data following a timecourse of Pol I.

B) Heatmap of Rpl/Rps genes showing a global decrease following various durations of iPol I. We noted in (A-B) there is some evidence of partial reduction in inhibition by CX-5461 by 8h; this may explain a lower induction of *Dux* compared to BMH-21 at this timepoint (Figure 4A) and could be due to reduced stability or feedback mechanisms. Conversely, 2C-specific gene upregulation is lower at 16-24h BMH-21 (Figure 3D), which may be due to apoptosis caused by abnormally high *Dux* expression.

C) Normalised log2 RNA-seq expression data showing upregulation of TEs belonging to the MERVL family at 8h following iPol I. CPM, counts per million.

D) Developmental progression rates past 2-cell before and 8h after the indicated treatments. P values, Chi-squared test, n >50 embryos from 4 experiments per condition.

**Figure S5** **Xie et al**

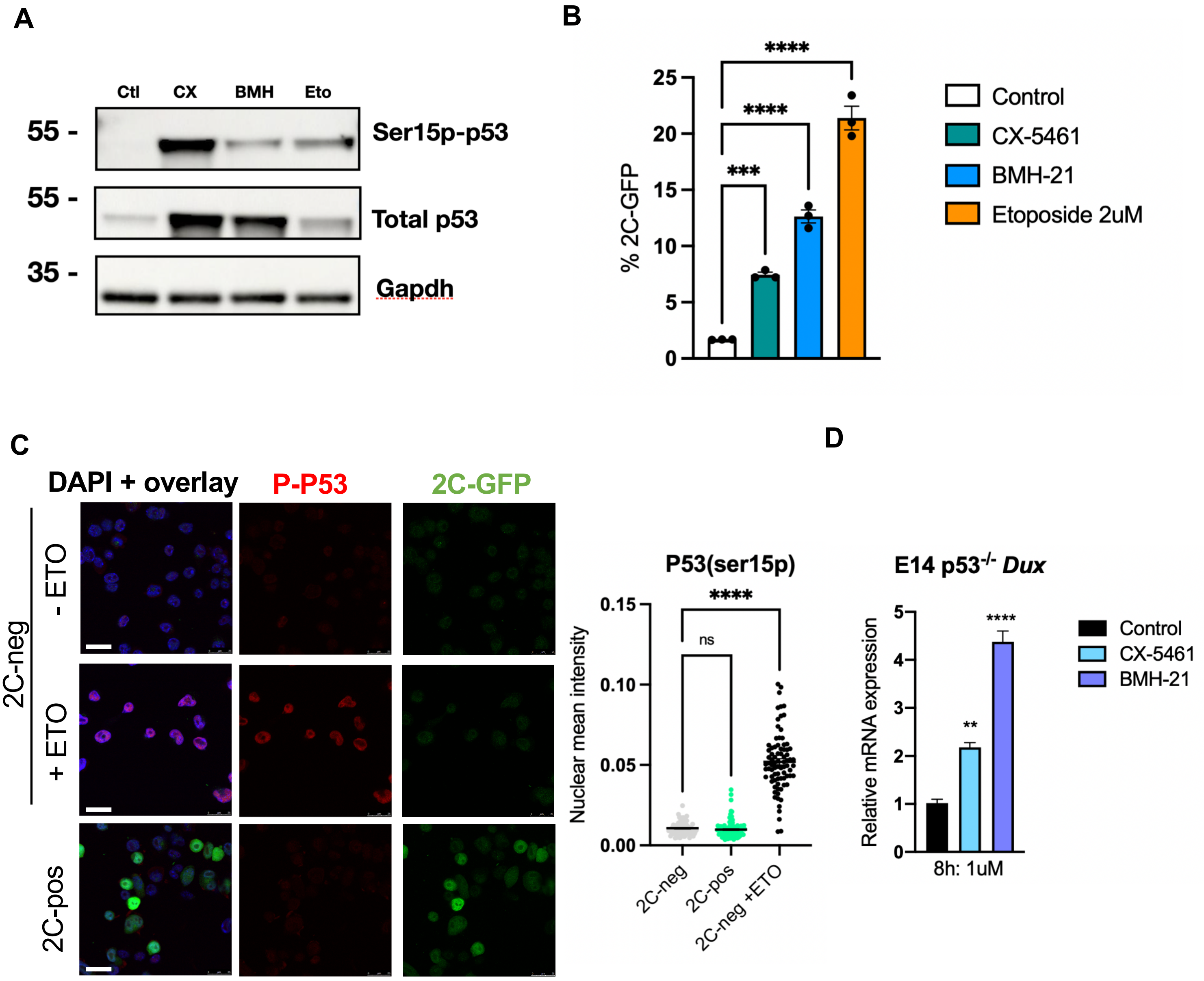


**Figure S5**. Mechanism of Dux and 2C activation upon iPol I

A) Western blots for total and phospho-p53 (Ser15p) in ESCs 24h after the indicated inhibitor treatments. Data are representative of 2 experiments.

B) % 2C-GFP+ cells as quantified by flow cytometry, 24h after treatment with the indicated inhibitors. Data are mean +/- s.e.m, n= 3 biological replicates.

C) Representative confocal microscopy images and quantification of phospho-p53 levels in 2C-negative and 2C-positive cells. Etoposide treatment was for 4h in 2C-negative cells as a positive control for p53 activation. Scale bars, 25 $\mu$m.

D) qRT-PCR analysis of *Dux* upregulation after iPol I in p53$^{-/-}$ ESCs, data are mean +/- s.e.m, 3 biological replicates.

All P values, one-way ANOVA with Dunnett multiple comparisons correction.

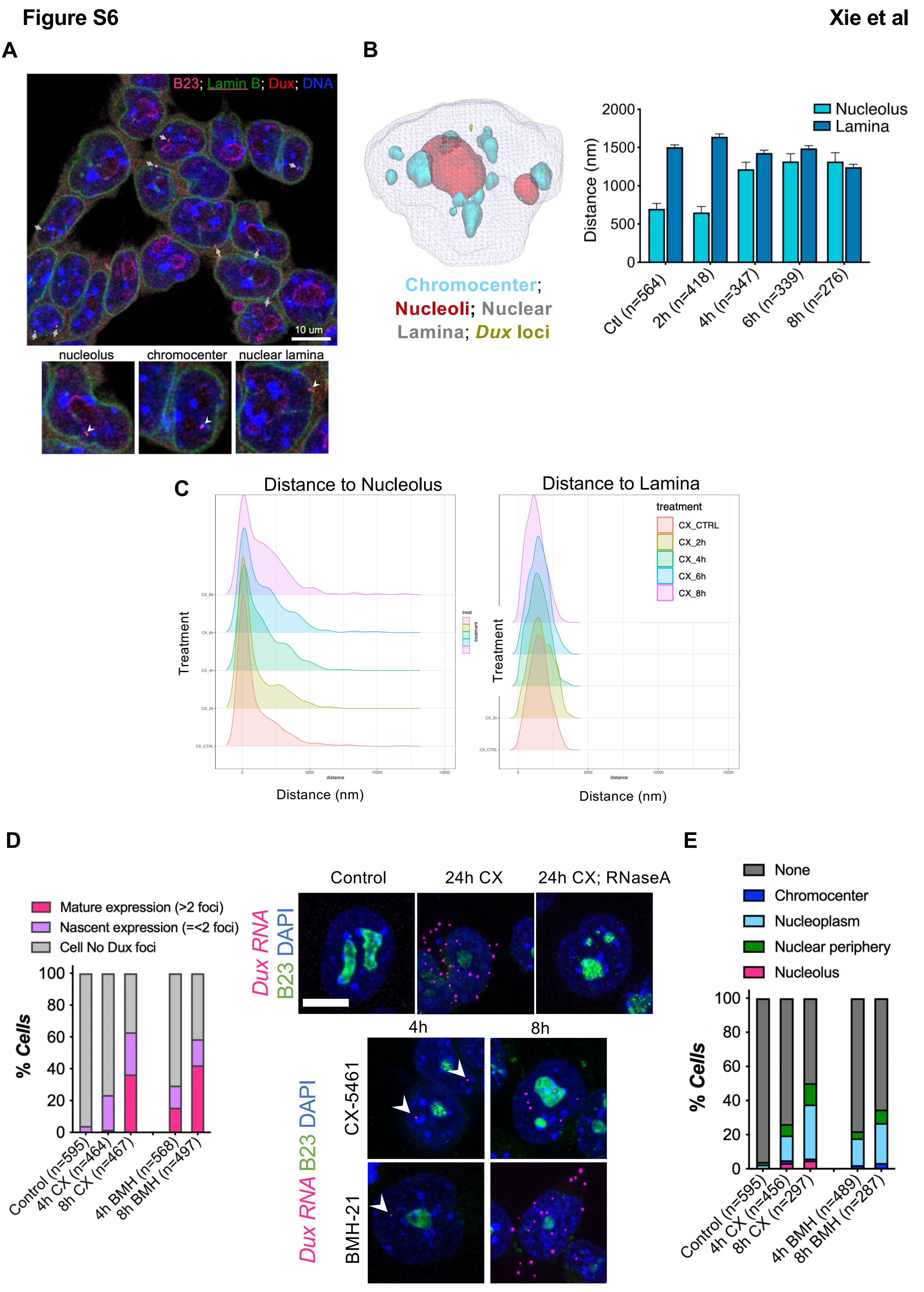

Figure S6
Xie et al
A
B23; Lamin B; Dux; DNA
10 um
nucleolus
chromocenter
nuclear lamina
B
Chromocenter; Nucleoli; Nuclear Lamina; Dux loci
Nucleolus
Lamina
Distance (nm)
2000
1500
1000
500
0
Ctl (n=564)
2h (n=418)
4h (n=347)
6h (n=339)
8h (n=276)
C
Distance to Nucleolus
Distance to Lamina
treatment
CX_CTRL
CX_2h
CX_4h
CX_6h
CX_8h
Treatment
Distance (nm)
D
Mature expression (>2 foci)
Nascent expression (=<2 foci)
Cell No Dux foci
% Cells
100
80
60
40
20
0
Control (n=595)
4h CX (n=464)
8h CX (n=467)
4h BMH (n=568)
8h BMH (n=497)
Control
24h CX
24h CX; RNaseA
Dux RNA B23 DAPI
4h
8h
CX-5461
BMH-21
E
None
Chromocenter
Nucleoplasm
Nuclear periphery
Nucleolus
% Cells
Control (n=595)
4h CX (n=456)
8h CX (n=297)
4h BMH (n=489)
8h BMH (n=287)

**Figure S6**. *Dux* movement upon iPol I.

A) Representative immuno-DNA-FISH for *Dux* DNA in ESCs alongside nucleolar and lamina markers with localization classification (below). *Dux* loci are marked by white arrowheads.

B) Example 3D reconstruction and identification of nuclear compartments and *Dux* loci in Imaris for calculation of 3D distances, and quantification of *Dux* movement from the nucleolus or lamina upon CX-5461 treatment. N=number of loci, data are mean +/- s.e.m for n= number of cells indicated.

C) Density plots of *Dux* distance (in nm) to the lamina or nucleolus upon CX-5461.

D) RNA FISH analysis of *Dux* expression in individual cells upon CX-5461 or BMH-21 treatment. Nascent expression in cells is defined as 2 or fewer nuclear foci, while mature expression is defined by >2 foci, n = number of cells scored. White arrowheads indicate nascent RNA foci. Scale bars, 10 $\mu$m.

E) Location of nascent RNA foci in ESCs upon iPol I, from scoring of cells with 2 or fewer *Dux* RNA foci, n = number of cells scored.

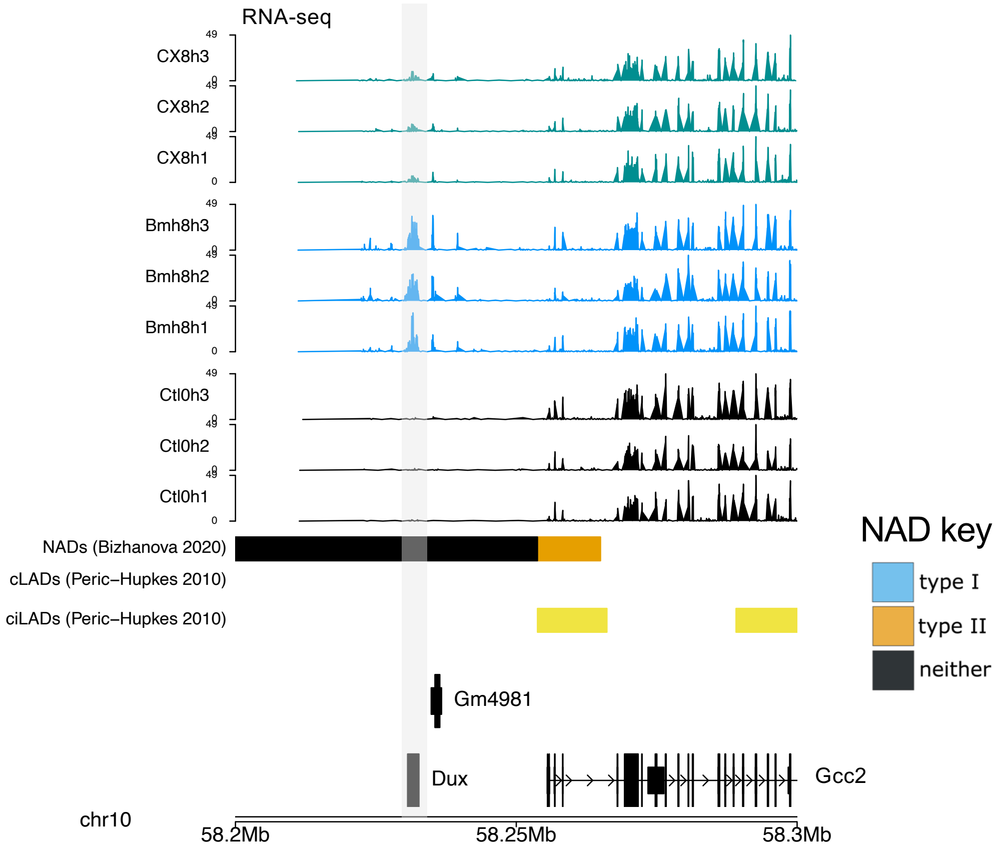


**Figure S7**. Browser screenshot of the indicated region of chr10 containing the *Dux* locus (highlighted) and neighbouring genes with overlayed RNA-seq tracks, confirming that *Dux* is situated in a NAD as defined in (Bizhanova et al., 2020). *Dux* is considered within a "neither" NAD, because its NAD does not overlap a cLAD or ciLAD - meaning in some cell types it is also lamina associated. This is consistent with some *Dux* loci positioning at the lamina in ESCs (Fig 5).
